# Evaluating the representational power of pre-trained DNA language models for regulatory genomics

**DOI:** 10.1101/2024.02.29.582810

**Authors:** Ziqi Tang, Nirali Somia, Yiyang Yu, Peter K Koo

## Abstract

The emergence of genomic language models (gLMs) offers an unsupervised approach to learning a wide diversity of *cis*-regulatory patterns in the non-coding genome without requiring labels of functional activity generated by wet-lab experiments. Previous evaluations have shown that pre-trained gLMs can be leveraged to improve predictive performance across a broad range of regulatory genomics tasks, albeit using relatively simple benchmark datasets and baseline models. Since the gLMs in these studies were tested upon fine-tuning their weights for each downstream task, determining whether gLM representations embody a foundational understanding of *cis*-regulatory biology remains an open question. Here we evaluate the representational power of pre-trained gLMs to predict and interpret cell-type-specific functional genomics data that span DNA and RNA regulation. Our findings suggest that probing the representations of pre-trained gLMs do not offer substantial advantages over conventional machine learning approaches that use one-hot encoded sequences. This work highlights a major gap with current gLMs, raising potential issues in conventional pre-training strategies for the non-coding genome.

## Introduction

Large language models (LLMs) have demonstrated remarkable capabilities in natural language processing^1–4^ and protein sequence analysis^5–8^. These LLMs, often termed “foundation models”, are trained through self-supervised learning to encode input data as contextual embeddings (also known as representations). The strength of pre-trained LLMs lies in the versatility of their embeddings, which can be leveraged for a broad spectrum of downstream predictive tasks. For instance, representations from pre-trained protein language models have been used to predict protein structures^9–11^, predict non-synonymous variant effects^12,13^, design novel protein sequences^14–16^, and study protein evolution^17,18^.

LLMs pre-trained on genomic DNA sequences offer a promising paradigm to accelerate our understanding of functional elements in the non-coding genome^19^. Genomic language models (gLMs) could, in principle, help to understand the complex coordination of transcription factors (TFs) to control the activity of *cis*-regulatory elements (CREs). They might also enable more accurate predictions of the functional consequences of non-coding mutations, which can help to prioritize disease-associated variants. Additionally, gLMs capable of learning *cis*-regulatory rules could become instrumental in designing novel regulatory sequences with desirable functional properties. They might also facilitate functional comparisons of non-coding sequences across different species, a task currently complicated due to substantial evolutionary drift in non-coding regions.

Recently, there has been a surge of pre-trained gLMs^20–49^. gLMs take as input DNA sequences that have undergone tokenization, an encoding scheme applied to either a single nucleotide or *k*-mer of nucleotides. Through self-supervised pre-training, the gLM learns a vector representation for each token in the DNA sequence via masked language modeling (MLM)^1^ or causal language modeling (CLM)^50^. In masked language modeling (MLM), a subset of input tokens undergo masking: most are replaced by a special `[MASK]` token, some by random tokens, and others left unchanged. The model learns to predict the original `[MASK]` tokens leveraging context from other unmasked positions, with random replacements introducing noise. Various masking strategies explore different granularities (words, phrases, entities) and approaches (permutations, sampling, importance-based selection) to enhance the self-supervised pre-training task’s effectiveness^51–55^. On the other hand, CLM is an autoregressive pre-training task with the goal of predicting the next token in a sequence given the previous tokens. Both language modeling objectives result in learning self-supervised representations of input sequences that capture information about individual tokens and the complex interrelationships with other tokens. The burden of learning biologically meaningful features is paid upfront during the pre-training. Afterward, the gLM’s representations can be leveraged for a broad spectrum of downstream prediction tasks as inputs to simpler models, bypassing the need to learn essential features for each task from scratch. In contrast, the conventional one-hot representation of DNA sequences treats each element independently, assigning an identical representation for the same nucleotide character irrespective of their position in the sequence or what context is nearby. Consequently, the responsibility of learning important patterns and their dependencies falls solely on the machine learning model being employed.

Current gLMs are composed of different choices for the tokenization, base architecture, language modeling objective, and pre-training data (Supplementary Table 1). *Tokenization* of DNA sequences is employed for either single nucleotide^20–22^ or *k*-mer of fixed size^23–25^ or a *k*-mer of variable sizes via byte-pair tokenization**^?^**^,27,33,56^, which aims to aggregate DNA in a manner that reduces the *k*-mer bias in the genome, a problem known as rare token imbalance. The *base architecture* is typically a stack of transformer layers^57^, with a vanilla multi-head self-attention^23–25,27–31^ or an efficient variant (e.g., flash attention^26,58^) or an exotic attention variant (e.g., sparse attention^32,33^). Alternatively, the base architecture has also been constructed with a stack of residual-connected dilated convolutional layers^20^ or selective state-space models, such as a Hyena^21,22,59^ or Mamba^43,46^. The *pre-training data* can vary significantly, encompassing the whole genome of a single species^20,24,32^ or the whole genomes across multiple species^23,25,26,28,33^ or focused only within specific regions of the genomes, such as the untranslated regions (UTRs)^29^, pre-mRNA^30^, non-coding RNA^60^, promoters^22^, coding regions^35,36,45^, non-coding RNA^39,60^, or conserved sites^34^.

Notably, Nucleotide-Transformer^23^ is a collection of BERT^1^-style models that consider non-overlapping *k*-mer tokenization and is pre-trained via MLM on either a single human genome, a collection of 3,202 human genomes from the 1000 Genomes Project^61^ alone or in combination with 850 genomes across diverse species. DNABERT2^26^ is also a BERT-style architecture but uses flash attention, considers byte-pair tokenization, and is trained via MLM on the genomes of 850 species. Genomic Pre-trained Network (GPN) is a convolution-based model with a stack of residual-connected dilated convolutions, uses single-nucleotide tokenization, and is trained via MLM on *Arabidopsis thaliana* genome and seven related species within the Brassicales order^20^. Similarly, HyenaDNA^21^ is a state-space model using Hyena layers and single-nucleotide tokenization and is trained via CLM on the human reference genome.

The utility of gLMs pre-trained on whole genomes for studying the non-coding genome has been limited. Previous benchmarks have largely considered gLMs that have been fine-tuned on each downstream task^23,24,26,30,39^. gLM fine-tuning involves adjusting the weights of all layers or through parameter efficient fine-tuning methods, such as LoRA (Low-Rank Adaptation)^26,62,63^, (hard or soft) prompt tuning^21,64^, and (IA)^323,65^. In each benchmark, a fine-tuned gLM has demonstrated improved predictions on a host of downstream prediction tasks, often based on the classification of functional elements, such as histone marks or promoter annotations. However, the chosen benchmarks do not reflect the complexity of *cis*-regulatory mechanisms observed in gene regulation, and the baseline models used in the comparisons often do not represent the state-of-the-art. Hence, the capabilities of gLMs in understanding the regulatory genome have yet to be demonstrated in a fair assessment.

However, fine-tuning makes it challenging to assess the contribution of the prior knowledge gained via pre-training on each downstream task. Moreover, benchmarks that do not fine-tune gLMs are limited in their downstream tasks^66–68^, relying on either binary classification of functional activity, which does not reflect the complexity of *cis*-regulatory biology^69,70^ or lack a more comprehensive set of benchmarking tasks. Thus, the extent to which existing gLMs pre-trained on whole genomes can genuinely serve as foundation models that can transfer their knowledge to predict and interpret functional genomics data without necessitating additional fine-tuning of the gLM weights.

**Figure 1.**
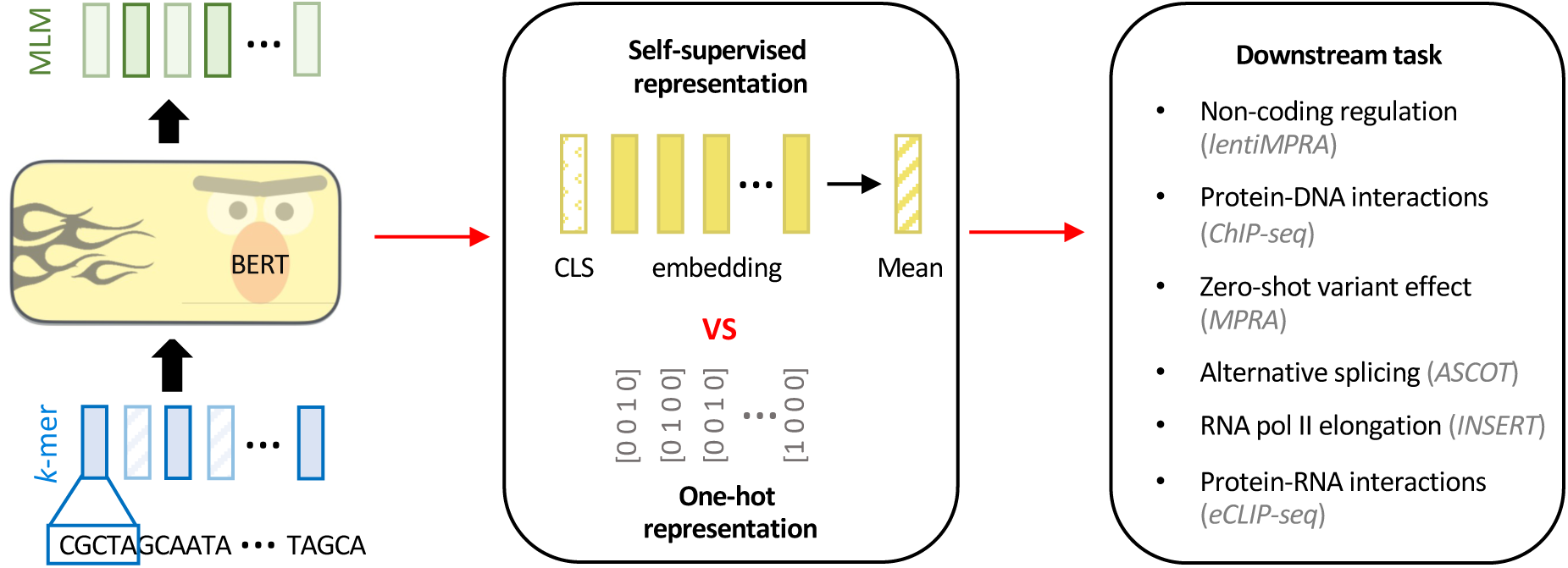
Experimental overview. Comparison of gLM embeddings versus one-hot representations for various functional genomics prediction tasks.

Here we perform a focused evaluation to assess the informativeness of learned representations of various gLMs pre-trained on whole genomes (without fine-tuning any existing layers) for six major functional genomics prediction tasks, which encompass different levels of *cis*-regulation complexity at DNA and RNA levels (see Fig. 1). In particular, we compared the predictive power via probing representations from pre-trained gLMs – namely Nucleotide-Transformer, DNABERT2, HyenaDNA, and a custom GPN pre-trained on the human reference genome – versus one-hot encoded DNA and representations acquired from a supervised “foundation model” pre-trained on a large corpus of functional genomics data. Our results suggest that current gLMs pre-trained on whole genomes do not provide noticeable advantages over conventional approaches to analyzing human functional genomics with deep learning using one-hot sequences. By contrast, supervised foundation models pre-trained on functional genomics data appear to encapsulate more relevant information and transfer better to other functional genomics data, albeit when the source pre-training tasks and the target tasks are closely aligned. Nevertheless, highly tuned supervised models trained from scratch using one-hot encoded sequences can achieve performance competitive with or better than pre-trained models across the datasets explored in this study. Our results suggest that current gLMs struggle to understand cell-type specific functional elements during pre-training and, therefore, fall short of recognition as a foundation model for the regulatory regions of the human genome.

## Results

### Task 1: Predicting cell-type specific regulatory activity from lentiMPRA data

Understanding the mechanisms that drive CRE activity is a major goal in functional genomics; it is challenging due to complex rules of cell-type-specific TF binding^71,72^. In the first task, we compared the performance of various machine learning models that consider different input representations of DNA sequences at predicting experimentally measured enhancer activity via lentiMPRA (lentiviral Massively Parallel Reporter Assay)^73^. Specifically, this task involves taking a 230 nucleotide (nt) DNA sequence as input, represented either as a gLM embedding or one-hot sequence, and predicting a scalar value that represents the CRE’s functional activity measured in a given cellular context via lentiMPRA (see Methods). This task enables a direct comparison in performance across matched downstream models for each sequence encoding scheme. By considering two cell types, namely HepG2 and K562, we can assess whether pre-trained gLM representations capture cell-type-specific CRE activity. While the original lentiMPRA study included the WCT11 cell type, we excluded it from our analysis due to the lack of correspondence with the cell types used in Task 3’s zero-shot single-nucleotide variant effect generalization.

For each gLM, we probed the embeddings from the penultimate layer using a linear model or multi-layer perceptron (MLP) based on the classification token (CLS) or the mean embedding, which is standard practice for harnessing sequence summarization of LLM embeddings. We also employed a baseline convolutional neural network (CNN) that analyzed the full embeddings of the penultimate layer as well as one-hot sequences for comparison (see Methods). We also considered embeddings from the penultimate layer of Sei^74^, a supervised foundation model pre-trained on 21,907 chromatin profiling datasets across over 1,300 cell lines and tissues. To assess the performance against a more sophisticated supervised model, we trained a ResidualBind^75^-like model (ResNet) using one-hot sequences. These choices provide a fair benchmark to assess whether embeddings from foundation models, acquired via unsupervised gLMs or supervised CNNs, are more informative for downstream models than naive one-hot sequences.

We found that a CNN trained on the whole sequence embedding led to improved performance over the linear or MLP models that analyzed CLS or mean embeddings (Fig. 2a). This suggests that summarized gLM representations lack sufficient information to predict cell-type-specific regulatory activity. In contrast, CNNs can build upon the full embeddings to better discriminate cell-type specific features. Moreover, the performance gap between MLPs and linear models suggests that the mapping between the pre-trained representations and the functional readouts of lentiMPRA data is highly non-linear. While a small-scale hyperparameter grid search showed comparable performance across different model capacity sizes (Supplementary Fig. 1), a more comprehensive architecture and hyperparameter search could potentially identify model settings that lead to further performance gains. However, for the scope of this study, we focused on simpler models, as is standard practice, to primarily assess the out-of-the-box utility of the learned gLM representations.

In a broader comparison, we also observed that CNNs trained using sequence embeddings from gLMs generally under-performed standard one-hot sequences, except our custom-trained GPN (Fig. 2b). Notably, the performance of all gLM-based representations was significantly lower than the supervised representations given by Sei and a LASSO regression baseline using features based on Enformer’s^76^ predictions, similar to Ref.^73^ (Supplementary Table 2). Due to differences in the data splits for Sei and Enformer, it is unclear to what extent data leakage might lead to performance inflation. Nevertheless, the ResNet model, which was trained from scratch using one-hot sequences from the LentiMPRA dataset, achieved the best performance (Fig. 2b). Although fine-tuning improved the predictive performance of the gLMs, achieving comparable performance as ResNet (Supplementary Table 2), these results suggest that pre-trained gLM embeddings may not provide beneficial context for CREs that cannot already be learned from one-hot sequences for the lentiMPRA dataset.

**Figure 2.**
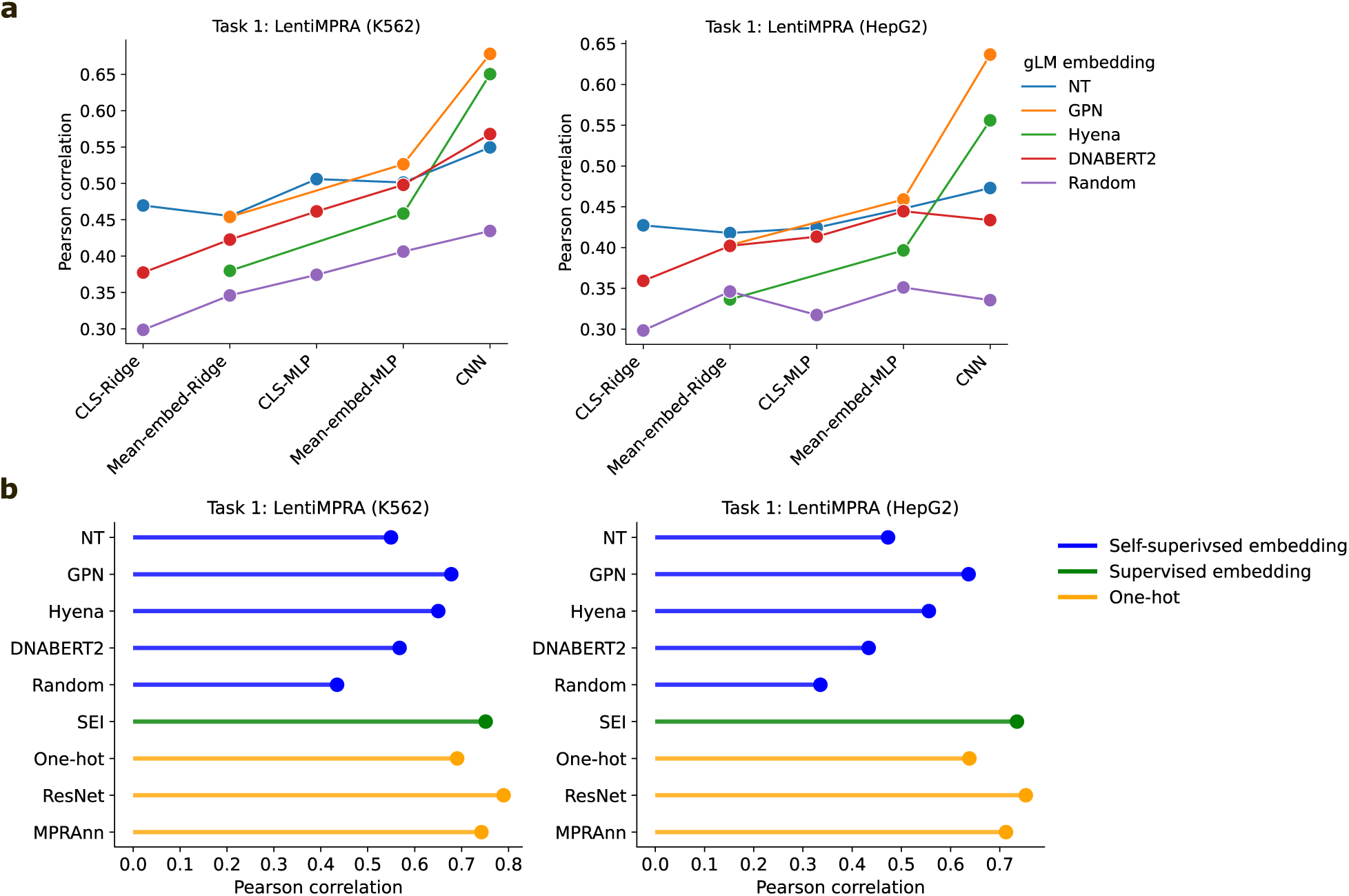
Performance comparison on cell-type-specific regulatory activity prediction tasks from lentiMPRA data. **a**, Comparison of predictive performance across various downstream machine learning models, including ridge regression and MLP using either the gLM’s CLS token or mean embedding, and a CNN trained using the full embedding of the penultimate layer of gLMs. **b**, Predictive performance using a baseline CNN trained using different gLM embedding inputs, one-hot sequences, or supervised embeddings from Sei. MPRAnn and ResNet represent the performance of more sophisticated models that are trained using one-hot sequences.

To control for the possibility that gLM embeddings from the penultimate layer may not be optimal, we performed the same analysis using embeddings from other layers of Nucleotide-Transformer. While some layers yielded modest improvements, particularly layer 10, the overall trends held and thus did not change the conclusions (Supplementary Fig. 2).

### Task 2: Predicting TF binding sites from ChIP-seq data

Since TF binding is a cell-type-specific phenomenon, but standard language modeling objectives are not cell-type aware, we surmised that the low performance of gLMs on the lentiMPRA prediction task may be due to losing information about key motifs during the pre-training. To test this hypothesis, we evaluated whether the gLM embeddings can predict TF binding sites measured via ChIP-seq (Chromatin Immuno-Precipitation sequencing^77^). Briefly, this task is framed as a binary classification where a model takes a 200 nt DNA sequence, either as a gLM embedding or a one-hot sequence, as input and predicts whether the sequence corresponds to a ChIP-seq peak. We consider ten ChIP-seq datasets spanning different TFs in GM12878 cells; a separate single-task model was trained for each TF (see Methods).

Evidently, CNNs trained using one-hot sequences modestly outperformed the whole embeddings from DNABERT2, HyenaDNA, and Nucleotide-Transformer. On the other hand, the custom GPN occasionally led to improved performance (Fig. 3). Since the TF binding tasks were included in the original pre-training of Sei, data leakage might lead to Sei’s inflated performance. Nevertheless, the modest performance differences across sequence encoding schemes, including the similar or better performance of one-hot encoding compared to gLM embeddings, suggest that the gLM embeddings are likely not actively encoding explicit information about transcription factor (TF) motifs. Rather, the embeddings appear to retain the essential sequence information necessary for downstream components like convolutional neural networks (CNNs) to learn TF binding patterns, akin to how CNNs can learn from one-hot encoded sequences that do not contain any inherent TF-related information.

**Figure 3.**
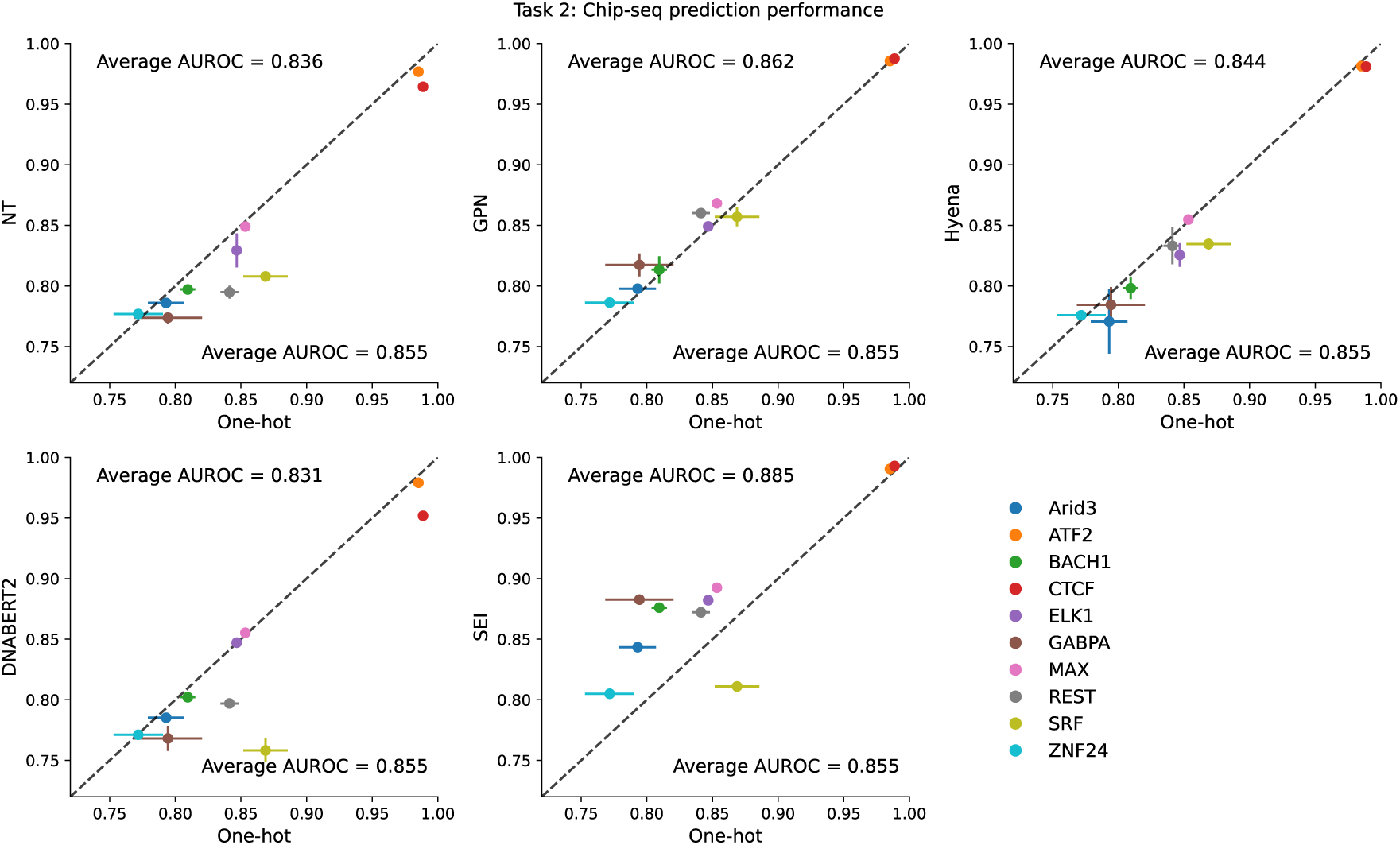
Performance comparison on TF binding prediction tasks from ChIP-seq data. Comparisons of CNNs trained using different gLM embeddings versus CNNs trained using one-hot sequences for 10 TF ChIP-seq datasets. Performance is measured by the average area-under the receiver-operating characteristic curve (AUROC) and error bars represent the standard deviation of the mean across 5 different random initializations. Average AUROC represents the average performance across all ChIP-seq datasets.

As a control experiment, we trained MLP or linear models using the CLS token of Nucleotide-Transformer. In this way, any information about motifs must be fully encoded in these summarized embeddings. We observed that CNNs trained on the whole embedding yielded substantially higher performance than an MLP trained using the CLS token (Supplementary Fig. 3a). However, the MLP still demonstrated proficiency in predicting TF binding overall. To further validate our findings and rule out the possibility of dataset biases creating a trivial prediction task, we also trained an MLP model on bag-of-dinucleotide frequencies. Indeed, the MLP based on dinucleotide frequencies yielded comparable performance to the CLS token (Supplementary Fig. 3a), except for CTCF, a protein that plays an important role in chromatin structure for all cell types. Together, these results suggest that gLMs do not appear to lose TF-related information in their embeddings, albeit only a slight information boost is gained regarding TF binding compared to low-level dinucleotide statistics. Nevertheless, downstream models that analyze conventional one-hot sequences can easily rectify any information deficiencies, leading to higher performances.

### Task 3: Zero-shot variant effect prediction with MPRA data

A major use case of highly accurate sequence-function models is their ability to predict the functional consequences of non-coding mutations^76^. In previous studies, Nucleotide-Transformer and GPN have demonstrated an ability to predict single-nucleotide variant effects, albeit as part of a binary classification task^20,23^. However, it is not intuitive how gLMs pre-trained on whole genomes could yield good zero-shot predictions of cell-type-specific variant effects in the non-coding region of human genomes since they are trained without any cell-type information. Thus, we assessed the ability of gLMs, specifically Nucleotide-Transformer, GPN, and HyenaDNA, to quantitatively predict single-nucleotide variant effects within CREs using saturation mutagenesis data measured via MPRAs (Massively Parallel Reporter Assay)^78^. This task involves calculating the zero-shot variant effect predictions of gLMs either by the cosine similarity of embedding vectors for the input sequence with mutant or wild-type allele (e.g., Nucleotide-Transformer and Hyena) or the log2-ratio of predicted variant and wild-type nucleotide via single-nucleotide masking (e.g., GPN). These variant effect scores are compared with experimentally measured variant effects according to the Pearson correlation coefficient (see Methods). This analysis includes MPRA measurements for three CREs in HepG2 cells and one CRE in K562 cells as part of the CAGI5 challenge^78,79^.

**Table 1.** Zero-shot variant effect generalization on CAGI5 dataset. The values represent the Pearson correlation between the variant effect predictions with experimental saturation mutagenesis values of a given CRE measured via MPRAs. Values are reported for a single CRE experiment for K562 and the average of three CRE experiments for HepG2.

| TRAINING TASK | MODEL | VARIANT EFFECT PREDICTION | HEPG2 | K562 |
| --- | --- | --- | --- | --- |
| SELF-SUPERVISED<br>PRE-TRAINING | NT (2B51000G) | COSINE DISTANCE | 0.125 | 0.007 |
|  | NT (2B5SPECIES) | COSINE DISTANCE | 0.112 | 0.135 |
|  | NT (500MHUMAN) | COSINE DISTANCE | 0.020 | 0.088 |
|  | NT (500M1000G) | COSINE DISTANCE | 0.041 | 0.068 |
|  | GPN (HUMAN) | LOG2-RATIO | 0.002 | 0.037 |
|  | HYENADNA | COSINE DISTANCE | 0.064 | 0.021 |
| LENTIMPRA-EMBEDDING | CNN-GPN | DIFFERENCE FROM WILD TYPE | 0.332 | 0.437 |
|  | CNN-NT | DIFFERENCE FROM WILD TYPE | 0.185 | 0.198 |
|  | CNN-SEI | DIFFERENCE FROM WILD TYPE | 0.579 | 0.701 |
| LENTIMPRA-ONE-HOT | CNN | DIFFERENCE FROM WILD TYPE | 0.324 | 0.365 |
|  | RESIDUALBIND | DIFFERENCE FROM WILD TYPE | 0.485 | 0.601 |
|  | MPRANN | DIFFERENCE FROM WILD TYPE | 0.381 | 0.437 |
| SUPERVISED ONE-HOT | SEI | COSINE DISTANCE | 0.545 | 0.641 |
|  | ENFORMER (DNASE) | DIFFERENCE FROM WILD TYPE | 0.510 | 0.685 |

We found that all tested gLMs (without fine-tuning) exhibited poor variant effect predictions in this quantitative zero-shot single-nucleotide generalization task (Table 1). These results extended to all Nucleotide-Transformer models^23^, including a 2.5 billion parameter BERT-based gLM trained on 3,202 diverse human genomes and 850 genomes from various species. On the other hand, CNNs trained using lentiMPRA data based on gLM embeddings yielded substantially better performance relative to their pre-trained counterparts (Table 1). Moreover, gLMs that were fine-tuned on the lentiMPRA data also yielded improved performance (Supplementary Table 3). In contrast, sophisticated supervised models trained using one-hot sequences, such as Enformer^76^, which is a state-of-the-art model trained with supervised learning on a wide variety of functional genomics data using one-hot sequences, and Sei yielded better performance than all CNNs trained using gLM representations. However, a CNN trained using Sei embeddings on the lentiMPRA dataset yielded the best overall performance. Together, these results highlight a major gap in the zero-shot variant effect performance of gLMs with the state-of-the-art.

### Task 4: Predicting alternative splicing from RNA-seq data

Previous studies demonstrated that Nucleotide-Transformer and GPN have learned properties related to gene definition and splice sites^20,23^. Thus, we surmised that gLMs pre-trained on whole genomes might be more beneficial for RNA regulation tasks. To investigate this, we tested the informativeness of gLM embeddings to predict mRNA alternative splicing quantified using RNA-seq (RNA-sequencing) from the ASCOT dataset^80^. Specifically, the prediction task takes as input two sequences – a sequence with 300 nt upstream of the splice acceptor and 100 nt downstream of the acceptor and a sequence with 100 nt upstream of the splice donor and 300 nt downstream of the donor – with the goal of predicting the percentage-spliced-in (PSI) across 56 tissues as a multi-task regression; a task introduced by MTSplice^81^. Similar to the DNA analysis, a baseline CNN was trained to take as input the full embeddings from gLMs or the embeddings of a pre-trained supervised model (see Methods).

Our results mirrored those seen for regulatory DNA, with embedding-based models largely under-performing compared to one-hot-based models (Fig. 4a). In contrast, Sei’s embeddings led to substantially lower performance than most gLM embeddings for this task. This is likely due to Sei’s pre-training focus on DNA-based functional genomics data, which leads to learning a set of DNA regulatory features that do not transfer well to RNA regulation. To test whether a more relevant set of features acquired through supervised learning could transfer better for RNA regulation, we trained a multi-task ResidualBind-like model to classify RNA-protein binding (RBP) sites from a large trove of eCLIP-seq data (see Methods). The task is to take 1,000 nt sequences as input and predict binding for 120 RBPs in K562 cells as a multi-task classification. Indeed, the embeddings from this RBP-trained supervised model led to substantially better performance than the gLM embeddings, except GPN, which yielded comparable results (Fig. 4a).

### Task 5: Predicting RNA pol II elongation potential from INSERT-seq data

Next, we performed a similar analysis for a prediction task that takes 173 nt RNA sequences as input and predicts RNA pol II elongation potential measured via INSERT-seq (INtegrated Sequences on Expression of RNA and Translation using high-throughput sequencing)^82^. The INSERT-seq dataset is modest in size, containing only 10,774 sequences. This small data regime may not provide sufficient examples to learn all relevant patterns using one-hot sequences. Training a large deep learning model on this dataset can easily lead to over-fitting. Thus, this task can help evaluate a scenario (i.e., the low data regime) where a baseline CNN that uses gLM embeddings might have an advantage over one-hot sequences.

**Figure 4.**
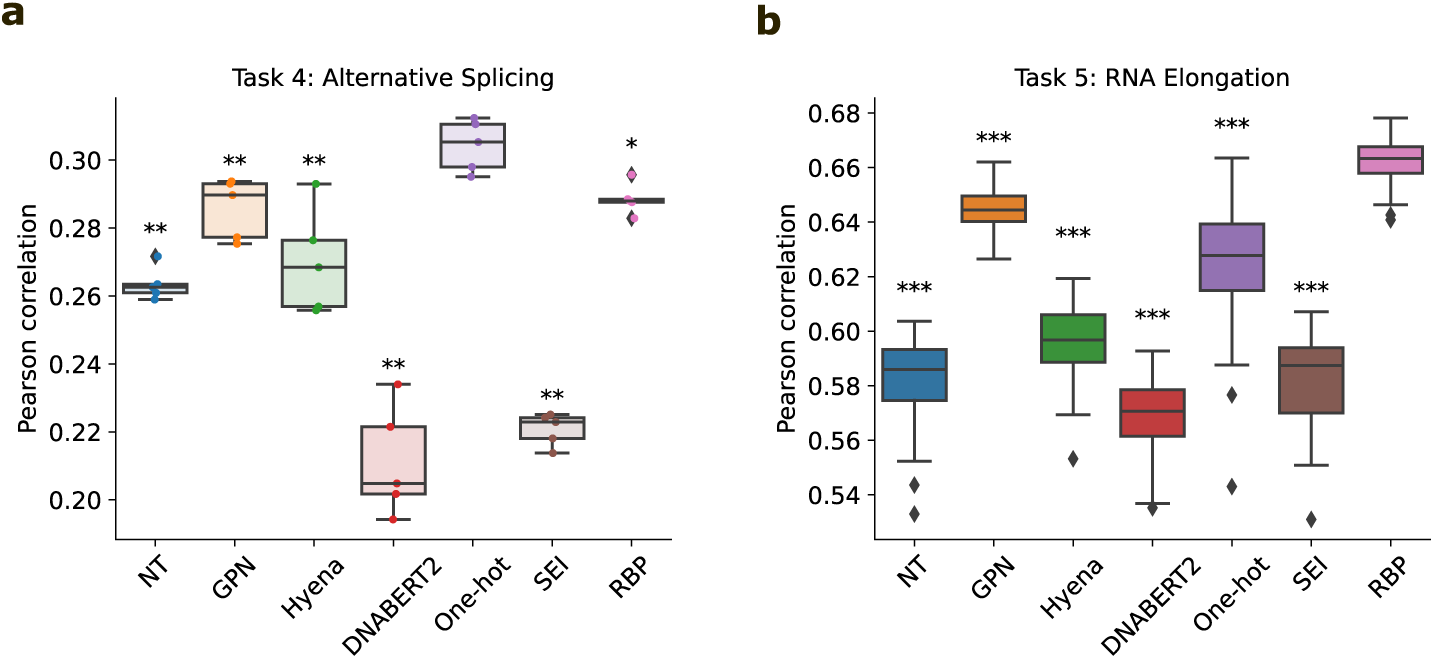
Performance comparison on RNA regulation tasks. **a**, Box-plots of the average Pearson correlation across tissues on test data for various models trained with different encoding schemes on an alternative splicing prediction task using MTSplice data. **b**, Box-plot of the Pearson correlation for various models trained with different encoding schemes on a RNA poll II elongation potential prediction task using INSERT-seq data. Box-plots show the first and third quartiles, central line is the median, and the whiskers show the range of data. Box-plots represent 5 different random initializations for **a** and 50 different random initializations for **b**. Statistical significance represents the Mann-Whitney U test with a *p* value *<* 0.05 (*^∗^*), *<* 0.01 (*^**^*), and *<* 0.001 (*^***^*).

Similarly, we found that the baseline CNNs trained using gLM embeddings yielded lower performance than one-hot RNA sequences, except for the custom GPN, which performed slightly better (Fig. 4b). Again, the CNN performance based on Sei’s supervised embeddings was worse, and the best-performing model was achieved using embeddings from the supervised multi-task model pre-trained to classify RBPs. These results highlight that generic pre-training strategies are not always beneficial; when carefully selecting pre-training tasks, one should consider which relevant features are needed to ensure more positive outcomes on downstream applications.

While the custom GPN was the only embedding that demonstrated improved performance over one-hot sequences, we hypothesized that further down-sampling of the training data could lead to situations where gLM embeddings become more beneficial than one-hot sequences. We systematically down-sampled both the alternative splicing and INSERT-seq datasets and retrained the same baseline CNNs using different input encoding schemes. Interestingly, the GPN embeddings consistently outperformed other embeddings (Supplementary Fig. 4). The improved performance by GPN suggests that gLMs may specialize more effectively in specific genomic regions. Specifically in this dataset, capturing 5’ splice sites is a critical feature^82^. Thus, understanding what features gLMs learn well can help to identify suitable downstream tasks for which they can thrive.

### Task 6: Predicting RNA-binding protein binding with eCLIP-seq data

RBPs are essential for various RNA processing stages, so next, we examined the ability of gLMs to predict RBP binding sites using eCLIP-seq (enhanced chromatin immunoprecipitation sequencing) datasets^83^. Briefly, the task involves taking 200 nt DNA sequences as input and predicting binary labels of whether the sequence corresponds to an eCLIP-seq peak or not (see Methods). Ten eCLIP-seq datasets spanning different RBPs were used in the evaluation. We trained a baseline CNN model using different sequence encoding schemes similar to previous tasks.

We found that CNNs trained using gLM embeddings performed slightly worse on average compared to the one-hot sequences (Fig. 5a), in agreement with the ChIP-seq results of Task 2. The narrow performance difference between models using gLM embeddings and one-hot sequences also indicates that RBP motif information is not lost in the gLM embeddings. In a similar control, we found that an MLP based on Nucleotide-Transformer’s CLS token led to slightly better performance than an MLP based on dinucleotide frequencies (Supplementary Fig. 3b). This supports that gLM embeddings encode beyond low-level sequence statistics in regulatory regions of RNA. Again, we found that Sei embeddings lead to a substantial decline in performance, further highlighting the importance of selecting appropriate pre-training tasks.

**Figure 5.**
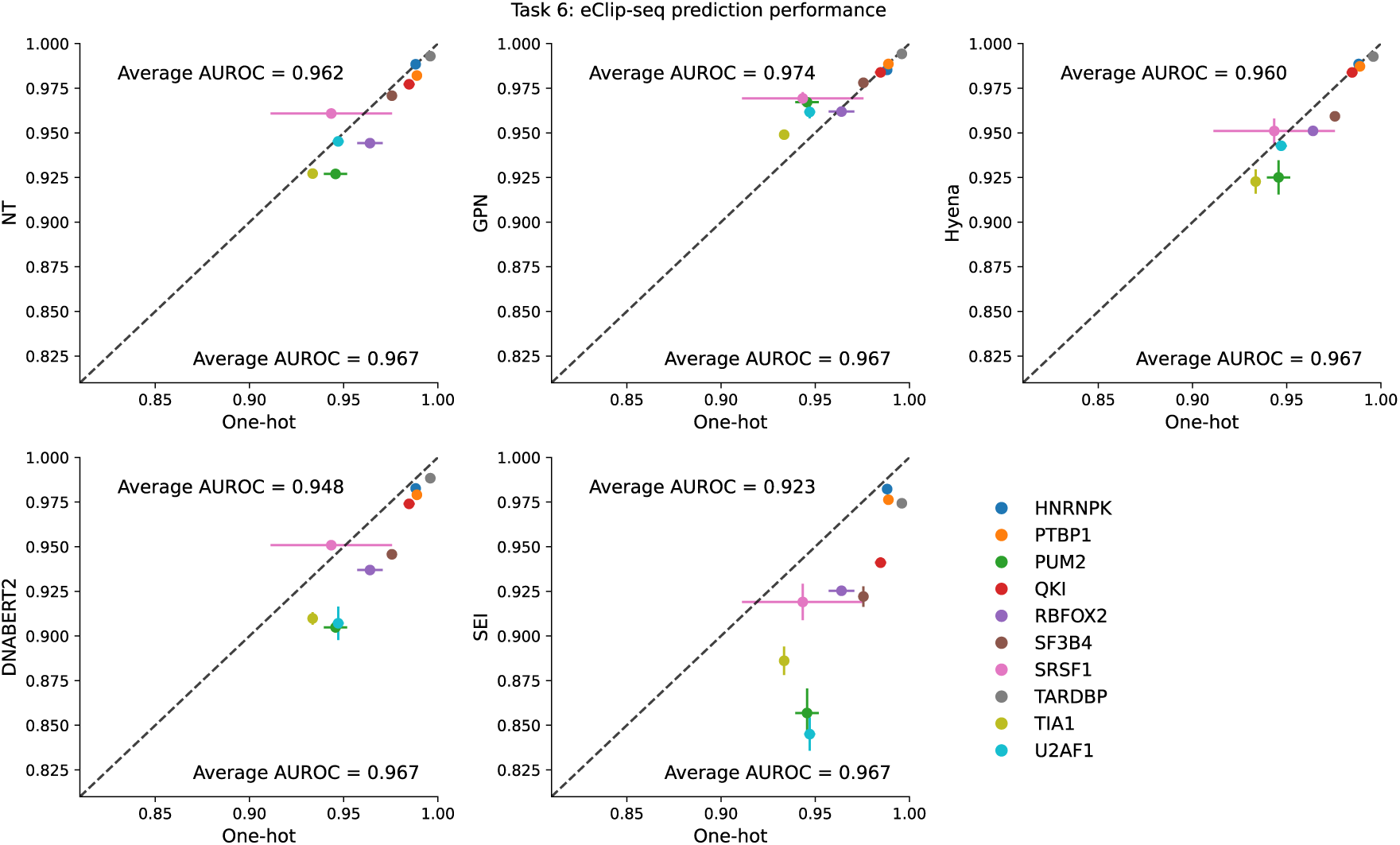
Performance comparison on RBP binding prediction tasks from eCLIP-seq data. Comparisons of CNNs trained using different gLM embeddings versus CNNs trained using one-hot sequences for 10 RBP eCLIP-seq datasets. Performance is measured by the average area-under the receiver-operating characteristic curve (AUROC) and error bars represent the standard deviation of the mean across 5 different random initializations. Average AUROC represents the average performance across all eCLIP-seq datasets.

### Uncovering cell-type-specific motifs learned by gLMs is challenging

As a follow-up, we performed an attribution analysis to identify motifs captured by gLMs. Attribution maps were generated for a given sequence by systematically masking one input token (i.e., a single nucleotide position for GPN and a non-overlapping *k*-mer for Nucleotide-Transformer) at a time and calculating the entropy over the predicted distribution of the masked token; ΔEntropy, which is the difference between the maximum entropy value across the whole sequence and the entropy values at each position, was used to identify positions that yielded informative nucleotides (see Methods). For comparison, we generated gradient-corrected Saliency Maps^84^ for a CNN trained using one-hot sequences. The analysis focused on lentiMPRA and CTCF ChIP-seq data to cover tasks from different systems with varying levels of complexity.

As expected, the attribution maps for pre-trained gLMs alone (i.e., not considering the downstream task) were difficult to interpret for both lentiMPRA (Fig. 6a) and ChIP-seq data (Supplementary Fig. 5a). The attribution maps did not reflect any known motifs, nor did they match any of the patterns captured in the CNN’s Saliency Maps. This disparity can arise if the probed locus is used across different cell types for multiple purposes. If cell-type-specific *cis*-regulatory patterns are projected onto a single DNA sequence, the overlapping set of motifs can lead to complex attribution maps that may not resemble distinct cell-type-specific motifs. Alternatively, the complex patterns that seem to span the length of the sequence could also reflect low-level sequence statistics that are memorized. Without ground truth, interpreting attribution maps remains challenging.

Next, we evaluated attribution maps generated by the downstream CNN that used gLM embeddings as input. Specifically, we scaled the gLM’s entropy-based attribution map with the maximum gradients at each position based on the downstream CNN (see Methods). Through a qualitative comparison, we noticed that the attribution maps generated by GPN appear to be visually aligned with Saliency Maps generated by the one-hot-trained CNN compared to Nucleotide-Transformer (Fig. 6a), even after accounting for the block-like structure which arises due to the *k*-mer tokenization. This trend was also observed for other loci (Supplementary Fig. 6).

To validate the importance of the putative binding sites identified via Saliency Maps for the one-hot-trained CNN, we employed global importance analysis (GIA)^75^. Specifically, we embedded the three annotated patterns into different dinucleotide-shuffled sequences, which serve as background sequences with low CRE activities, and measured the effect of including the patterns on model predictions. Indeed, GIA shows that the motif patterns identified by Saliency Maps for the one-hot-trained CNN are more or less sufficient to explain model predictions (Fig. 6b).

We then quantified the correlation between the attribution maps generated by the one-hot-trained CNN and the gLM-based attribution maps. We found that attribution maps generated by pre-trained gLM are not well-aligned with each other, nor are the attribution maps generated by the one-hot-trained CNN (Fig. 6c, Supplementary Fig. 5b). By contrast, attribution maps generated by CNNs trained with gLM embeddings led to improved alignment between their attribution maps and with one-hot-trained CNNs. These results suggest that the gLMs learn non-overlapping features during pre-training, but a downstream model can still use them to build cell-type-specific motifs (that are better aligned with motifs learned by one-hot-trained CNNs). Together, the attribution maps given by pre-trained gLMs seem to visually capture a more diffuse set of patterns, which speculatively reflect low-level statistics of genomic sequences. Downstream models, like CNNs, use these seemingly uninformative gLM embeddings (especially from GPN) to build cell-type-specific regulatory features relevant for downstream prediction tasks.

**Figure 6.**
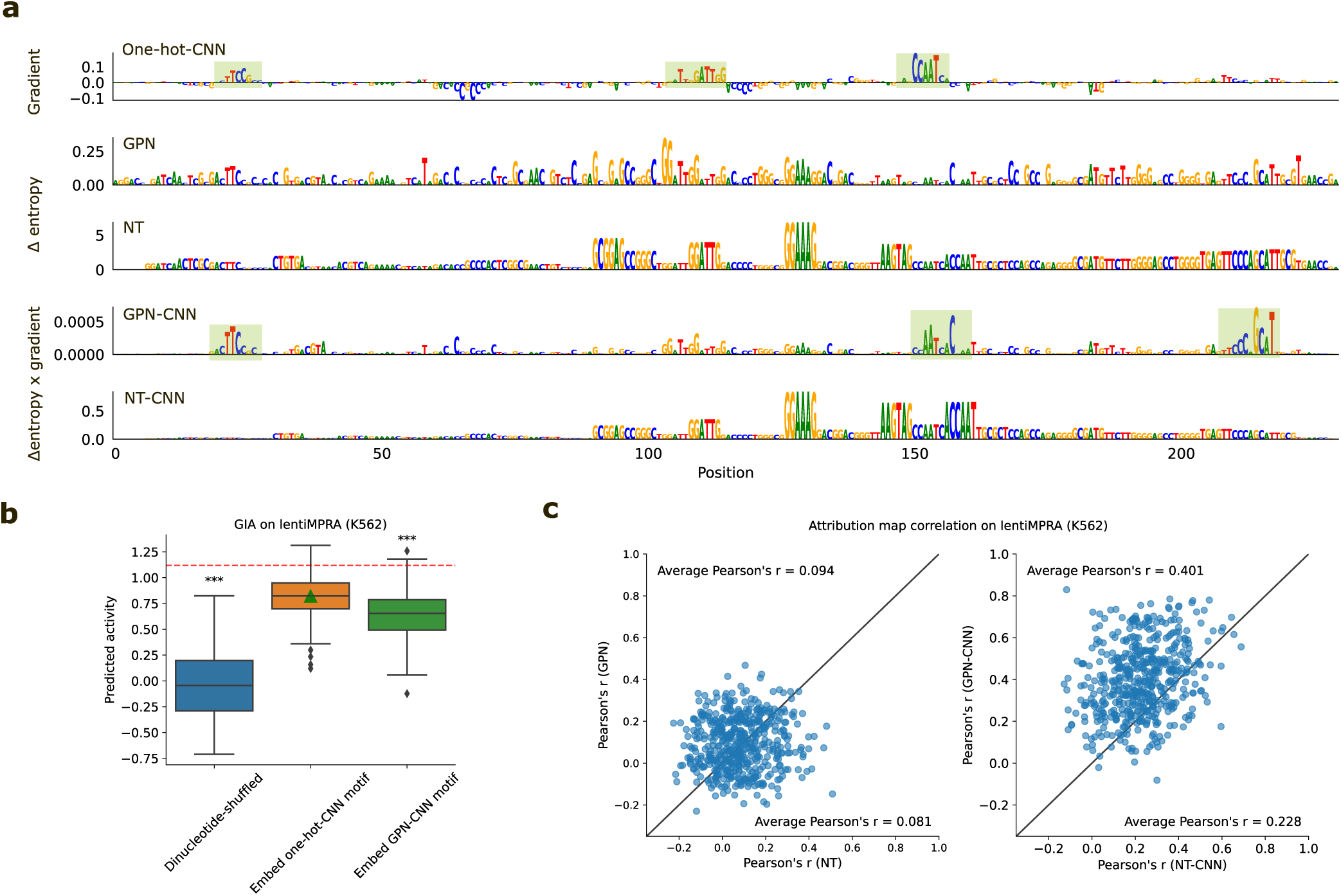
Attribution analysis comparison for sequences from the lentiMPRA dataset. **a**, Representative example of attribution maps for a regulatory sequence. Attribution maps include (top to bottom): the gradient-times-input of a one-hot-trained CNN; the delta entropy of predicted nucleotides via single-nucleotide masking from a pre-trained GPN; the delta entropy of predicted nucleotides via single-nucleotide masking from a pre-trained Nucleotide-Transformer; the gradient of a CNN-trained using GPN embeddings multiplied by the delta entropy of predicted nucleotides via single-nucleotide masking from a pre-trained GPN; and the gradient of a CNN-trained using Nucleotide-Transformer embeddings multiplied by the delta entropy of predicted nucleotides via single-nucleotide masking from a pre-trained Nucleotide-Transformer. **b**, Box-plot of the predicted activity for 300 dinucleotide-shuffled sequences from **a**, dinuc-shuffled sequences with the annotated patterns from the Saliency Map of the one-hot-trained CNN, and dinuc-shuffled sequences with the annotated patterns from the CNN trained using GPN embeddings (GPN-CNN). Green triangle represents the global importance analysis value. Red dashed line represents the prediction of the wild type sequence according to the one-hot-trained CNN. Box-plots show the first and third quartiles, central line is the median, and the whiskers show the range of data. **c**, Scatter plot comparison of the attribution map correlations for different pre-trained gLMs (left) and CNNs trained using gLM embeddings (right). Attribution map correlations reflect the Pearson correlation coefficient between the attribution map generated by the gLM-based attribution method with the Saliency Map generated by a one-hot-trained CNN. Each dot represents a different sequence in the lentiMPRA dataset (N=500).

## Discussion

To assess the transferability of knowledge acquired during pre-training for current genome language models, we evaluated the predictive power of pre-trained representations from four gLMs on whole genomes (without fine-tuning) across six functional genomics prediction tasks with appropriate baselines for comparison. We found that within *cis*-regulatory elements, representations from pre-trained gLM provide little to no advantage compared to standard one-hot sequences. On a relative basis, we found that GPN, a convolution-based LLM, yielded slightly more informative representations in the non-coding genome compared to highly parameterized BERT-style LLMs. This suggests that stronger inductive biases toward learning relevant features in the model architecture might improve gLMs, albeit modestly.

Notably, we elected not to fine-tune weights of the gLM on each downstream task, which is how gLMs have been previously benchmarked^23,24,26,30,39^. While gLM performance undoubtedly improves with fine-tuning, the scope of this study was to gauge the knowledge of *cis*-regulatory biology learned by gLMs during pre-training. The poor performance observed in this study suggests that cell-type-specific *cis*-regulatory mechanisms are predominantly learned during fine-tuning. Our results suggest that the benefit of pre-training gLMs appears to be initializations that are pre-loaded with just a little more information than low-level statistical properties for non-coding genomic sequences. Further research is needed to understand how biological knowledge is refined from pre-training to fine-tuning.

In previous studies, pre-trained gLMs have found some success by focusing on specific regions of the genome during pre-training or working with simpler organisms with compact genomes^28,34,85^. For instance, a BERT-based LLM trained strictly in the coding genome can provide more context than only considering amino-acids with protein language modeling (e.g., codon usage)^35,36,45^. However, our evaluation shows that extending the pre-training task across the whole genome struggles to capture meaningful representations in the non-coding genome.

The performance gap may be due to differences in the structure of the coding regions versus the non-coding regions. To elaborate, protein sequences have a clear start and end with low-level grammars (i.e., secondary structures) and high-level grammars (i.e., protein domains) shared throughout most globular proteins, with structures conserved across species. On the other hand, the non-coding genome contains a variety of short sequence motifs that vary broadly in binding affinities and are sparsely located in seemingly random DNA, with usage and rules that vary across loci and cell types. Few non-coding elements exhibit deep conservation that is typical in proteins. The differing selection pressures in the non-coding regions lead to loss of synteny, which makes it difficult to study sequence and functional conservation. Thus, treating each nucleotide position equally, whether informative or uninformative, makes this a challenging language modeling task. In the non-coding genome, this is tantamount to expecting the LLM to predict predominantly random nucleotides, which, by definition, can only be achieved via memorization. Hence, this may explain why gLMs have also found greater utility in learning *cis*-regulatory features in simpler organisms with compact genomes, such as bacteria^40,85,86^, arabidopsis^20^, or yeast^28^, which have substantially reduced junk DNA^87–89^.

We note that recent supervised foundation models, such as Enformer^90^ and Borzoi^91^, may serve as better examples of supervised foundation models but their scale makes it difficult to probe their transferrability to the small-scale sequences utilized in this study. Their large inputs, which are hundreds of kilobases long, require either substantial zero-padding, which introduces a substantial covariate shift, or sequence context marginalization tricks^69,75,92^, which is computationally expensive. Moreover, the non-uniformity in data splits makes any direct comparison challenging due to potential inflated performance from data leakage. In future evaluations, we plan to include more foundation models, including Enformer, Borzoi, and new gLMs that emerge, focusing on a broader set of chromatin-based functional genomics prediction tasks.

A major benefit of gLMs is their lack of reliance on labels generated from wet-lab experiments during training, allowing them to learn a broader set of patterns. However, our results suggest that gLMs have yet to learn a foundational set of *cis*-regulatory features in the non-coding genome of humans that can be harnessed via probing in prediction tasks across cell types. By contrast, supervised deep learning models trained on large troves of functional genomics data in a multitask setting can learn discriminative features related to *cis*-regulatory mechanisms in the non-coding genome^70,90,91,93–96^. However, the representations learned by these models are biased towards the experiments they are trained on, which are predominantly generated within a few cell lines. Hence, their generalization capabilities to other cell types remain limited.

Current gLMs require fine-tuning to achieve comparable performance as an optimized supervised model trained using one-hot sequences. While this highlights a weakness of the foundational knowledge learned during pre-training, it can still be considered beneficial. This is because gLMs can be conveniently fine-tuned on a wide variety of downstream tasks and achieve competitive performance without the challenge of optimizing sensitive hyperparameters of a supervised model on each downstream task.

Evaluating what gLMs have learned through predictive modeling remains an endless endeavor. A more efficient approach can be achieved through model interpretation of the gLMs, which should help to understand the alignment between gLMs and prior biological knowledge. Our preliminary analysis of attribution maps was inconclusive, highlighting the need for a more in-depth understanding of what gLMs are learning from pre-training. Further development can build upon the initial progress^97–99^ towards more meaningful domain-inspired model interpretation tools to bridge this gap.

Looking forward, it remains an open question whether LLMs will bring the same revolution in human genomics as in other fields. The current trends in scaling gLMs (via larger models and considering broader sequence contexts^21,32^) might only produce incremental gains, albeit achieved inefficiently according to scaling laws^100^, as the availability of diverse and informative genomics data is a major limiting factor. It remains unclear whether continued scaling of the gLMs pre-trained with standard language modeling objectives (i.e., MLM or CLM) will eventually lead to realizing emergent capabilities, such as learning cell-type-specific *cis*-regulatory biology in the non-coding genome. The amount of genetic variation required to capture the full complexity of the human genome may be simply too great, as a single genome encodes for the spatio-temporal regulation of all cell types. Additional information, such as functional genomics data, is likely needed during the pre-training for gLMs to become proficient in characterizing cell-type specific functional elements. Even protein language models trained solely on amino-acid sequences can learn conservation and protein structure elements and yet struggle to generalize well to a wide diversity of functional tasks^101^. At the least, a separate language modeling objective for different regions in the genome to account for the high entropy in the non-coding regions is needed. Due to the high upfront costs to train gLMs with the lack of reciprocal performance gains on downstream tasks, gLMs will likely require a more focused, domain-inspired revelation in pre-training objectives to achieve the esteemed “foundation” status for the non-coding genome.

## Methods

### Pre-trained language models

#### Nucleotide-Transformer

Nucleotide-Transformer consists of multiple BERT-based language models with 2 different model sizes (i.e., 500 million and 2.5 billion parameters) and trained on various sets of genome sequences: human reference genome, 1000 genomes project, and 850 genomes from several species. Details of the tokenizer, model structure, and training procedure can be found in the original paper^23^. We acquired weights for each Nucleotide-Transformer model from the official GitHub repository. In this analysis we mostly used representations from NT2.5B-1000G, except for the zero-shot variant effect generalization analysis, which considered all Nucleotide-Transformer models. Since Nucleotide-Transformer models allow flexible input sizes, no padding was necessary for any evaluation tasks.

#### Custom GPN

The GPN model is a convolutional neural network that was originally trained on Arabidopsis genome sequences via masked language modeling with an input size of 512 nucleotides^20^. It consists of 25 convolutional blocks, where each convolutional block includes a dilated convolutional layer followed by a feed-forward layer, connected by intermediate residual connections and layer normalization. The dilation rate for each convolutional layer cycles with increasing exponentially by factors of 2, from 1 to 32. The embedding dimension was kept fixed at 512 throughout the layers. For our custom GPN (human) model, we created training datasets using the human reference genome (hg38^102^). The genome was split into contigs and filtered for a minimum length of 512 nucleotides, with chromosome 8 held out as test set. During training, 15% of the nucleotide positions were masked and the model is tasked to predict the nucleotide probabilities for each masked location. The model was trained for 2 million steps with a constant learning rate of 0.001 using ADAM^103^.

#### HyenaDNA

The HyenaDNA model is a gLM pre-trained on the human reference genome, with context lengths up to 1 million tokens at the single nucleotide-resolution^21^. Architecturally, it adopts a decoder-only, sequence-to-sequence configuration, organized into a succession of blocks each encompassing a Hyena operator^59^, followed by a feed-forward neural network. The model weights and representation extraction code was acquired through the Hugging Face repository^104^. For all experiments in this study, we used the “hyenadna-tiny-1k-seqlen-d256” model due to the sequence length limitation of the functional genomics datasets.

#### DNABERT2

DNABERT2, a second generation version of the original DNABERT model^24^, is constructed on the BERT architecture, comprising 12 Transformer blocks. In this new iteration, the authors improved the model by replacing learned positional embeddings with Attention with Linear Biases (ALiBi) and utilizing Flash Attention to increase computation and memory efficiency^26^. In the context of this study, analyses were done with the representations generated by the last Transformer block. The model was acquired through the Hugging Face repository, using the “DNABERT-2-117M” model.

### Pre-trained supervised models

#### Sei

The Sei model is composed of three sequential modules: (1) a convolutional network with dual linear and nonlinear paths; (2) residual dilated convolution layers; (3) spatial basis function transformation and output layers. Sei was trained to take as input 4 kb length sequences and predict 21,907 TF binding, histone marks and DNA accessibility from peak data of *cis*-regulatory profiles. For this study, we extracted our representations after the spline basis function transformation, before inputting into fully connected layers. The pre-trained Sei model was acquired through zenodo from the original study^74^.

#### RBP

Our custom RBP model was trained using eCLIP-seq^83^ data of 120 RBPs in K562 from ENCODE^105^. The dataset was organized into a multi-task binary classification format. The model has a ResidualBind-like structure:

1. 1D convolution (96 filters, size 19, batch-norm, exponential) dropout (0.1)
2. Dilated residual block^106^

convolution (96 filters, size 3, batch-norm, ReLU)
dropout (0.1)
convolution (96 filters, size 3, batch-norm, dilation rate 2)
dropout (0.1)
convolution (196 filters, size 3, batch-norm, dilation rate 4)
dropout (0.1)
skip connection to input
ReLU activation
max-pooling (size 10)
dropout(0.1)
3. 1D convolution (192 filters, size 7, batch-norm, ReLU)

dropout (0.1)
global average-pooling
4. flatten
5. fully-connected (512 units, batch-norm, ReLU)

dropout (0.5)
6. output layer (120 units, sigmoid)

#### Enformer

A LASSO regression was fit to the lentiMPRA data based on predictions from Enformer^76^. Each sequence was padded to a target length of 196,608 base pairs using zero-padding on both ends. The padded sequences were then passed through Enformer to generate predictions. We extracted the predictions corresponding to the central bin (bin=448). Lasso regression was employed using scikit-learn. The training data was split into training (80%) and validation (20%) sets. LassoCV with 5-fold cross-validation was employed to select the optimal regularization parameter (alpha) from 200 candidates. The model was trained with a maximum of 20,000 iterations and a tolerance of 1e-2. Performance was evaluated using the Pearson correlation coefficient on the test set. Unlike previously^73^, the predictions were based on a single model trained on one fold and only considering the forward strand.

### Data

#### lentiMPRA

The lentiMPRA dataset for K562 and HepG2 cell lines was acquired from the Supplementary Tables in Ref.^73^. The HepG2 library consists of 139,984 sequences, each 230 nucleotides long, and the K562 library contains 226,253 sequences. Each sequence is paired with a target scalar value that represents the transcriptional activity. Each cell line was treated independently as a single-task regression. For each dataset, we randomly split the training, validation, and test sets according to the fractions 0.7, 0.1, 0.2, respectively. Unlike the original study, we treated reverse-complement sequences separately; they were not aggregated or augmented during test time. The results represent the performance over a single fold.

#### CAGI dataset

The CAGI5 challenge dataset^78^ was used to evaluate the performance of the models on zero-shot single-nucleotide variant effect generalization as following the same procedure as Ref.^69^. We only considered MPRA experiments in HepG2 (LDLT, SORT1, F9) and K562 (PKLR). We extracted 230 nucleotide sequences from the reference genome centered on each regulatory region of interest. Alternative alleles are then substituted correspondingly to construct the CAGI test sequences. Pearson correlation was calculated between the varient effect scores by the model and experimentally measured effect size per experiment. For HepG2 performances, we report the average Pearson’s r across the three experiments.

#### ChIP-seq

Ten transcription factor (TF) chromatin immunoprecipitation sequencing (ChIP-seq) datasets were acquired from the zenodo repository of Ref.^84^. The prediction task is a binary classification of whether 200nt input DNA sequences are associated with a ChIP-seq peak (positive label) versus sequences from DNase I hypersensitive sites from the same cell type (i.e., GM12878) that do not overlap with any ChIP-seq peaks (negative label). The number of negative sequences were randomly down-sampled to exactly match the number of positive sequences to ensure balanced classes. The dataset was split randomly into training, validation, and test set according to the fractions 0.7, 0.1, and 0.2, respectively.

#### Alternative splicing data

Data was acquired from direct correspondence with the authors of Ref.^81^ Briefly, 61,823 cassette exons from ASCOT was split into a training, validation, and test set. The training set consisted of 38,028 exons from chromosome 4, 6, 8, 10-23, and the sex chromosomes. The 11,955 exons from chromosome 1, 7, and 9 were used as the validation set, and the remaining 11,840 exons were used as the test set (chromosomes 2, 3, and 5). Models are evaluated based on their performance on the test set. The prediction task takes as input two sequences – a sequence with 300 nt upstream of the acceptor and 100 nt downstream of the acceptor and a sequence with 100 nt upstream of the donor and 300 nt downstream of the donor – and the goal is to predict PSI across 56 tissues as a multi-task regression.

#### INSERT-seq

INSERT-seq data was obtained from Ref.^82^. INSERT-seq measures the impact of transcribed sequences on the RNA polymerase II elongation potential and expression in mouse embryonic stem cells. 11,417 insert sequences of length 173nt long were used as inputs and the goal is to predict the totalRNA output, which measures the relative abundance in RNA relative to genomic DNA, as a regression task. Training, validation, and test sets were split according to the fractions 0.8, 0.1, and 0.1, resulting in 9,131, 1,149, and 1,137 sequences, respectively.

#### eCLIP datasets

The *in vivo* eCLIP-based datasets were downloaded from the ENCODE. For each RBP experiment, the bed narrowPeaks (two replicates) and the bam file for the corresponding mock inputs experiment were downloaded. For each replicate, we removed peaks with a signal value less than 1 and a log-*p*-value greater than 3. Using bedtools, the remaining peaks that share at least one nucleotide across the two replicates were selected as positive peaks. A correlation filter across the replicates was applied: 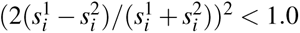, where 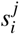 represent the signal value for the *i*th peak in replicate *j*. The median peak size was about 50 nt with a positive tail that exceeded 200 nt in some cases. Positive sequences were generated by extracting 200 nucleotide sequences about the center position of the peak coordinates. Sequences with undefined nucleotides were filtered out. Negative peaks were generated by employing Piranha peak caller on the bam file of the mock inputs with a bin size of 20 and a *p*-value threshold of 0.01. We then removed negative peaks which overlap with any unfiltered peaks from each replicate. Negative peaks were generated by extracting 200 nt sequences about the center position of the remaining negative peak coordinates. Because the negative peaks usually had more entries compared to the positive peaks, we randomly selected a similar number of negative peaks as positive peaks. All sequences were given a corresponding label 1 for sequences which contain a positive peak and 0 for sequences which contain a negative peak. All sequences were then randomly split into a training set, validation set, and test set according to the fractions 0.7, 0.1, and 0.2, respectively.

### Models for downstream tasks

#### Linear models

Linear models with L2 regularization (i.e., Ridge) serve as the baseline, representing a simple downstream model. The inputs of the model were based on the embeddings of the CLS token or the average embedding across sequences for Nucleotide-Transformer models. For regression and classification tasks, the linear model was a linear regression or logistic regression, respectively. The strength of the L2 regularization was set to 1e-3.

#### MLP

A multi-layer perceptron model was used to train on CLS token embeddings or the average embedding across sequences for Nucleotide-Transformer models. The model is constructed by two fully connected blocks. The first block includes a fully-connected layer with 512 units and ReLU ativation, followed by batch-normalization and a dropout rate of 0.5. The second block consists of a fully-connected layer with 256 units and the same activation, batch-normalization, and dropout layers. The model was trained on lentiMPRA dataset with Adam optimizer, learning rate of 0.0001, mean-squared error loss function, learning rate decay with a patience of 5 epochs and a decay factor of 0.2, and early stopping patience of 10 epochs.

#### MPRAnn for lentiMPRA

MPRAnn is a convolutional based model with a total of 4 convolutional and 3 dense layers trained on the lentiMPRA dataset. It takes 230 nt one-hot encoded sequences including the adapters as input to predict the mean log2(RNA/DNA) values from forward and reverse strands. We augmented the batches using the reverse-complement of the 200 nt target sequence, while keeping the two 15 bp adapters fixed. To fit the model, we used a learning rate of 0.001, an early stopping criterion with patience of 10 on 100 epochs, and the Adam optimizer with a mean square error loss function. Model structure and training parameters obtained from Github directory of original publication^73^.

#### Baseline CNN for lentiMPRA

We designed a baseline CNN model with the following structure:

1. batch-norm (optional)
2. 1D convolution (196 filters, size 1) (optional)
3. 1D convolution (196 filters, size 7, batch-norm, exponential)

dropout (0.2)
max-pooling (size 5)
4. 1D convolution (256 filters, size 7, batch-norm, ReLU)

dropout (0.2)
max-pooling (size 4)
5. flatten
6. fully-connected (512 units, batch-norm, ReLU)

dropout (0.5)
7. fully-connected (256 units, batch-norm, ReLU)

dropout (0.5)
8. output layer (1 unit, linear)

CNN models were trained with Adam optimizer, mean-squared error loss function, learning rate of 0.0001 with a learning rate decay patience of 5 epochs with a decay rate of 0.2, and early stopping with patience of 10 epochs for both one-hot sequence and language model embedding-based training on the lentiMPRA dataset. For one-hot sequences, batch-norm and the convolution with kernel 1 were not employed.

**ResidualBind for lentiMPRA.** We designed the ResidualBind model by adding a dilated residual block after the first convolutional layer of the baseline CNN model, according to:

1. 1D convolution (196 filters, size 19, batch-norm, Silu)

dropout (0.2)
2. Dilated residual block

convolution (196 filters, size 3, batch-norm, Silu) dropout (0.1)
convolution (196 filters, size 3, batch-norm, Silu, dilation rate 2) dropout (0.1)
convolution (196 filters, size 3, batch-norm, Silu, dilation rate 4) dropout (0.1)
convolution (196 filters, size 3, batch-norm, Silu, dilation rate 8) dropout (0.1)
convolution (196 filters, size 3, batch-norm, Silu, dilation rate 16) dropout (0.1)
convolution (196 filters, size 3, batch-norm, dilation rate 32) skip connection to input
Silu activation
max-pooling (size 5)
dropout(0.2)
3. 1D convolution (256 filters, size 7, batch-norm, Silu)

dropout (0.2)
max-pooling (size 5)
4. fully-connected (256 units, batch-norm, Silu)

dropout (0.5)
average-poolint (size 2)
5. flatten
6. fully-connected (256 units, batch-norm, Silu)

dropout (0.5)
7. output layer (1 unit, linear)

ResidualBind was trained with Adam optimizer, mean-squared error loss function, learning rate of 0.001 with a learning rate decay patience of 5 epochs with a decay rate of 0.2, and early stopping with patience of 10 epochs.

#### Baseline CNN for ChIP-seq and CLIP-seq

We designed a baseline CNN model with the following structure:

1. batch-norm (optional)
2. 1D convolution (512 filters, size 1) (optional)
3. 1D convolution (64 filters, size 7, batch-norm, ReLU)

max-pooling (size 4)
dropout (0.2)
4. 1D convolution (96 filters, size 5, batch-norm, ReLU)

max-pooling (size 4)
dropout (0.2)
5. 1D convolution (128 filters, size 5, batch-norm, ReLU)

max-pooling (size 2)
dropout (0.2)

1. flatten
2. fully-connected (256 units, batch-norm, ReLU)

dropout (0.5)
3. output layer (1 unit, linear)

CNN models were trained with Adam optimizer, binary cross-entropy loss function, learning rate of 0.001 with a learning rate decay patience of 5 epochs with a decay rate of 0.2, and early stopping with patience of 10 epochs for both one-hot sequence and language model embedding-based training on the lentiMPRA dataset. For one-hot sequences, batch-norm and the convolution with kernel 1 were not employed.

#### Insert-seq model

For the RNA pol II elongation potential dataset, we developed a residual convolutional network structure and used it for all embedding and one-hot-based models. The model was trained using mean square error loss function, Adam optimizer, learning rate of 0.0001, learning rate decay patience of 5 epochs with a decay rate of 0.2, and early stopping patience of 10 epochs.

1. convolution(48 filters, size 1) (optional)
2. convolution (96 filters, size 19, batch-norm, exponential)

dropout (0.1)
3. dilated residual block

convolution (96 filters, size 3, batch-norm, ReLU)
dropout (0.1)
convolution (96 filters, size 3, batch-norm, dilation rate 2)
dropout (0.1)
convolution (96 filters, size 3, batch-norm, dilation rate 4)
skip connection to block input
ReLU activation
max-pooling (size 10)
dropout(0.1)
4. convolution (128 filters, size 7, batch-norm, ReLU)

global average-pooling
dropout (0.1)
5. fully-connected layer (128 units, ReLU)

dropout (0.5)
6. output layer (1 unit, linear)

CNN models were trained with Adam optimizer, mean-squared error loss function, learning rate of 0.0001 with a learning rate decay patience of 5 epochs with a decay rate of 0.2, and early stopping with patience of 10 epochs for both one-hot sequence and language model embedding-based training on the lentiMPRA dataset. For one-hot sequences, the convolution with kernel 1 was not employed.

### Zero-shot variant effect prediction methods

For Nucleotide-Transformer, we derived the zero-shot predictions using cosine similarity as suggested in the original study^23^. For each variant, we passed the sequences with the centered reference allele and the alternative allele through the model to extract embeddings. The cosine similarity between the two complete sequence embeddings was calculated and used as the zero-shot score. A negative correlation is expected between the score and effect size. Since this distance-based zero-shot score only reflects the magnitude, not the direction, of function change, we calculated the Pearson correlation using the absolute value of the effect size.

For GPN, we followed a similar procedure as the original study^20^. First, we input sequences with the center variant loci masked and acquired the predicted allele probabilities for the masked loci. Then, we calculate the zero-shot prediction score as the log-likelihood ratio between the alternate and reference alleles. Again, since the likelihood ratio doesn’t reflect the direction of function change associated with the variants, we calculated the correlation score using the absolute value of effect size.

Finally, for the embedding-based and one-hot based models, we used the difference in predictions between the alternative and reference allele sequence as the zero-shot prediction score. For Enformer, we use the cell-type agnostic approach of averaging the effect size across all DNase-seq tracks. To reduce predictions to scalars, we summed across the profile predictions.

### Attribution methods

For CNN models, the attribution analysis was based on grad-times-input with saliency maps. The gradients of the prediction were calculated with respect to the input sequence to yield an L x A map, where L is the length of the sequence and A is 4 (one for each nucleotide). By subtracting the position-wise average saliency scores from this map and then multiplying by the one-hot encoded sequence, the method isolates the sensitivity of each observed nucleotide at every position, enhancing interpretability by pinpointing nucleotide-specific contributions to predictions.

For gLMs, the analysis involved sequentially masking each token of the input sequence and predicting the probability of the masked token by the model. The entropy of the probability distribution for each position was computed to quantify the information content represented by the gLM. Given that lower entropy signifies a higher information level, the saliency score was derived as the difference between the maximum entropy value and the entropy at each position, ensuring that a higher saliency score reflects greater information retention.

Sequence logos were visualized using Logomaker^107^.

### Global importance analysis

Global importance analysis was carried out according to Ref.^75^. A example sequence was selected from the LentiMPRA (K562) dataset. We sampled 300 dinucletoide shuffled versions of the sequence to be used as background sequences. The shuffling aims to preserve the dinucleotide frequency while destroying any coherent patterns. The LentiMPRA trained One-Hot-CNN models’ predictions for the shuffled sequences are considered to be the baseline for predicted CRE activity. The top three positive motif patters identified separately in the One-hot-CNN and GPN-CNN saliency maps (Fig. 6c) were inserted into the corresponding position of the shuffled sequences, creating two experiment sequences sets. The One-Hot-CNN model was used to make predictions for the motif embedded sequences. The difference in prediction for the three sets of sequences reflect the global importance of these motif patterns to the CNN model.

### Model Fine-tuning on lentiMPRA Data

We fine-tuned DNABERT2, HyenaDNA, and Nucleotide Transformer on lentiMPRA data derived from HepG2 and K562 cell lines. Each model was adapted for sequence regression, predicting a single continuous value corresponding to CRE activity.

#### Nucleotide Transformer

We used the 500M parameter version pre-trained on 1000 Genomes data. Fine-tuning employed the LoRA technique, applied to the query and value matrices of the self-attention mechanism. The LoRA rank was set to 1, with a scaling factor of 32 and dropout rate of 0.1. An AdamW optimizer with a learning rate of 5e-4 was used, training for 1000 steps or 2 epochs (whichever came first) with a batch size of 64.

#### HyenaDNA

The pre-trained `hyenadna-tiny-1k-seqlen` model was used for fine-tuning. The model performed a mean pool of the penultimate representation across sequence length and then used a linear layer to transform the pooled representation to a single output. We employed a character-level tokenizer for DNA bases (A, C, G, T, N), with a maximum sequence length of 230 tokens and left-side padding. Training used PyTorch with an AdamW optimizer, learning rate of 6e-4, and weight decay of 0.1 for 100 epochs with a batch size of 256. The final model checkpoint after 100 epochs was used for evaluation.

#### DNABERT2

We utilized the `DNABERT-2-117M` model, fine-tuning it using the Hugging Face Transformers library. An AdamW optimizer with a learning rate of 3*e&−* 5 was used, training for 5 epochs with batch sizes of 8 and 16 for training and evaluation, respectively.

For all models, mean squared error was used as the loss function. The lentiMPRA datasets for both cell lines were preprocessed and stored in HDF5 format. Model performance was evaluated using mean squared error, Pearson correlation coefficient, and Spearman correlation coefficient. All experiments were conducted using CUDA-enabled GPUs, with the best-performing model for each combination selected based on the lowest validation loss.

## Data Availability

Processed data and model weights can be found at: https://doi.org/10.5281/zenodo.11583224. Datasets include lentiMPRA (Task 1), ChIP-seq (Task 2), MPRAs for zero-shot single-nucleotide generalization (Task 3), alternative splicing (Task 4), INSERT-seq (Task 5), and eCLIP-seq (Task 6).

## Code Availability

Open-source code to reproduce this study is available on GitHub (https://github.com/amberT15/LLM_eval) as well as ref.**^?^**.

## Acknowledgements

The authors would like to thank Evan Seitz for providing Enformer variant effect predictions and other members of the Koo Lab for helpful comments on the manuscript. Research reported in this publication was supported in part by the National Human Genome Research Institute of the National Institutes of Health under Award Number R01HG012131 and the National Institute Of General Medical Sciences of the National Institutes of Health under Award Number R01GM149921. ZT was also supported by the Elisabeth Sloan Livingston Fellowship. Funding for NS was provided by the National Institute of Health Postbaccalaureate Research Experience Program at Cold Spring Harbor Laboratory (NIGMS PREP award R25GM144246). This work was performed with assistance from the US National Institutes of Health Grant S10OD028632-01.

## Author contributions

ZT and PKK conceived of the method and designed the experiments. ZT developed code, ran the experiments, and analyzed the majority of the results. YY and NS performed the fine-tuning analysis for gLMs for Task 1 and Task 3. NS performed the Enformer analysis for Task 1. ZT and PKK interpreted the results and contributed to writing the paper.

## Competing interests

Nothing to declare.

---

**Supplementary Table 1.** Summary of existing gLM architecture and training choices.

| Name | Base Model | Training Data | Pre-training Task | Tokenization | Finetuned for evaluations |
| --- | --- | --- | --- | --- | --- |
| GPN [20] | CNN | 8 plant genomes | MLM | Base-res | No |
| GPN-MSA [34] | BERT | Conserved regions in 100 vertebrates MSA (PhastCons) | MLM | Base-res | No |
| Nucleotide Transformer [23] | BERT | 1000 Genomes, Human genome and multi-species genomes | MLM | Non-overlapping k-mer | Yes |
| DNABERT [24] | BERT | Human genome | MLM | Overlapping k-mer | Yes |
| DNABERT2 [26] | BERT-Flash attention | Human genome and multi-species genomes | MLM | Byte-pair encoding | Yes |
| BigBird [32] | Sparse attention | Human genome | MLM+CLM | Byte-pair encoding | Yes |
| hyenaDNA [21] | Hyena | Human genome | CLM | Base-res | Yes |
| OmniNA [31] | LLaMA | Multi-species and text annotations | CLM (multimodal) | Byte-pair encoding | Yes |
| GENA-LM [33] | BERT | T2T, 1000 Genomes, and multi-species genomes | MLM | Byte-pair encoding | Yes |
| LOGO [38] | ALBERT | Human genome | MLM | Non-overlapping k-mer | Yes |
| DNA-GPT [25] | GPT | Genome from 9 species | CLM, GC-content, and sequence order | Non-overlapping k-mer | Yes |
| GROVER [27] | BERT | Human genome | MLM | Byte-pair encoding | Yes |
| UTR-LM [29] | BERT | 5' UTR sequences (Ensembl) | MLM and supervised (secondary structure and minimum free energy) | Base-res | Yes |
| SpliceBERT [30] | BERT | pre-mRNA from 72 vertebrates | MLM | Base-res | Yes |
| RegLM [22] | Hyena | Yeast promoter sequences | CLM | Base-res | Yes |
| RNA-FM [39] | BERT | RNAcentral (promoters) | MLM | Base-res | Yes |
| Species-aware DNA LM [28] | BERT | UTR of 1500 yeast genomes | MLM | Base-res | No |
| FloraBERT [41] | BERT | Multi-species plant genome | MLM | Byte-pair encoding | Yes |
| RandomMask [42] | BERT | Human genome | MLM (expanding size) | Overlapping k-mer | Yes |
| GenSLMs[40] | BERT | BV-BRC dataset | Contrastive loss | Codon | Yes |
| Evo [85] | StripedHyena | prokaryotic genomes and plasmids | CLM | Base-res | Both |
| Mamba [43] | Mamba | human genome | CLM | Base-res | Both |
| Caduceus [46] | Bi-directional Mamba | prokaryotic genomes and plasmids | MLM | Base-res | Yes |
| PlantCaduceus [47] | SSM (Bi-directional Mamba) | 16 Angiosperm genomes | MLM | Nucleotide | Yes |
| AgroNT [48] | Transformer (BERT) | 48 plant genomes | MLM | Non-overlapping k-mer | Yes |
| Epigenomic BERT [49] | Transformer (BERT) | Human genome, epigenetic state information (IDEAS) | MLM | Non-overlapping k-mer | Yes |
| RiNALMo [60] | Transformer (BERT) | non-coding RNA | MLM | Nucleotide | Both |

**Supplementary Table 2.**
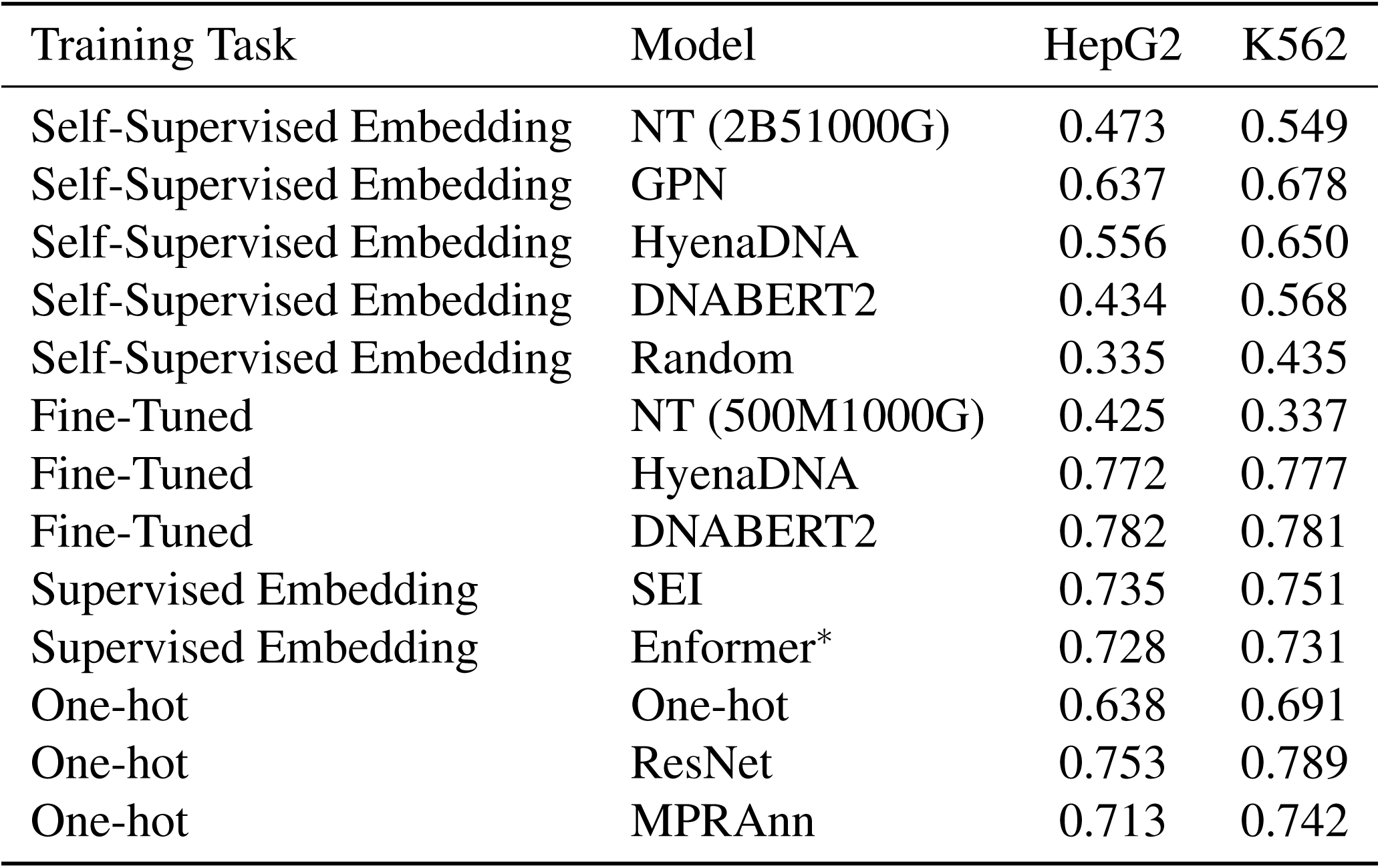
Performance comparison on cell-type-specific regulatory activity prediction tasks from lentiMPRA data. Predictive performance using a baseline CNN trained using different gLM embedding inputs, one-hot sequences, or supervised embeddings from Sei. MPRAnn and ResNet represent the performance of more sophisticated models that are trained using one-hot sequences. The performance of NT, Hyena, and DNABERT2 fine-tuned directly on the lentiMPRA data is also shown. Note that NT fine-tuning is different from the self-supervised embedding due to limited resources for fine-tuning. Enformer*^*^* represents a LASSO regression – i.e., linear probing based on Enformer’s predictions, instead of a downstream CNN trained on the full embeddings like SEI.

**Supplementary Table 3.** Expanded zero-shot variant effect generalization on CAGI5 dataset. The values represent the Pearson correlation between the variant effect predictions with experimental saturation mutagenesis values of a given CRE measured via MPRAs. Values are reported for a single CRE experiment for K562 and the average of three CRE experiments for HepG2. Notably, this table includes the performance based on NT, HyenaDNA and DNABERT2 when fine-tuned on lentiMPRA data.

| Training Task | Model | Variant Effect Prediction | HepG2 | K562 |
| --- | --- | --- | --- | --- |
| Self-Supervised Pre-Training | NT (2B51000G) | cosine distance | 0.125 | 0.007 |
| Self-Supervised Pre-Training | NT (2B5Species) | cosine distance | 0.112 | 0.135 |
| Self-Supervised Pre-Training | NT (500MHuman) | cosine distance | 0.020 | 0.088 |
| Self-Supervised Pre-Training | NT (500M1000G) | cosine distance | 0.041 | 0.068 |
| Self-Supervised Pre-Training | GPN (Human) | log2-ratio | 0.002 | 0.037 |
| Self-Supervised Pre-Training | HyenaDNA | cosine distance | 0.064 | 0.021 |
| Fine-Tuned | NT (500M1000G) | difference from wild type | 0.155 | 0.104 |
| Fine-Tuned | HyenaDNA | difference from wild type | 0.286 | 0.315 |
| Fine-Tuned | DNABERT2 | difference from wild type | 0.357 | 0.430 |
| LentiMPRA-Embedding | CNN-GPN | difference from wild type | 0.377 | 0.457 |
| LentiMPRA-Embedding | CNN-NT (2B51000G) | difference from wild type | 0.137 | 0.240 |
| LentiMPRA-Embedding | CNN-SEI | difference from wild type | 0.559 | 0.719 |
| LentiMPRA-one-hot | CNN | difference from wild type | 0.313 | 0.426 |
| LentiMPRA-one-hot | Residualbind | difference from wild type | 0.486 | 0.551 |
| LentiMPRA-one-hot | MPRAnn | difference from wild type | 0.301 | 0.369 |
| Supervised one-hot | SEI | cosine distance | 0.545 | 0.641 |
| Supervised one-hot | Enformer (DNase) | difference from wild type | 0.510 | 0.685 |

**Supplementary Figure 1.**
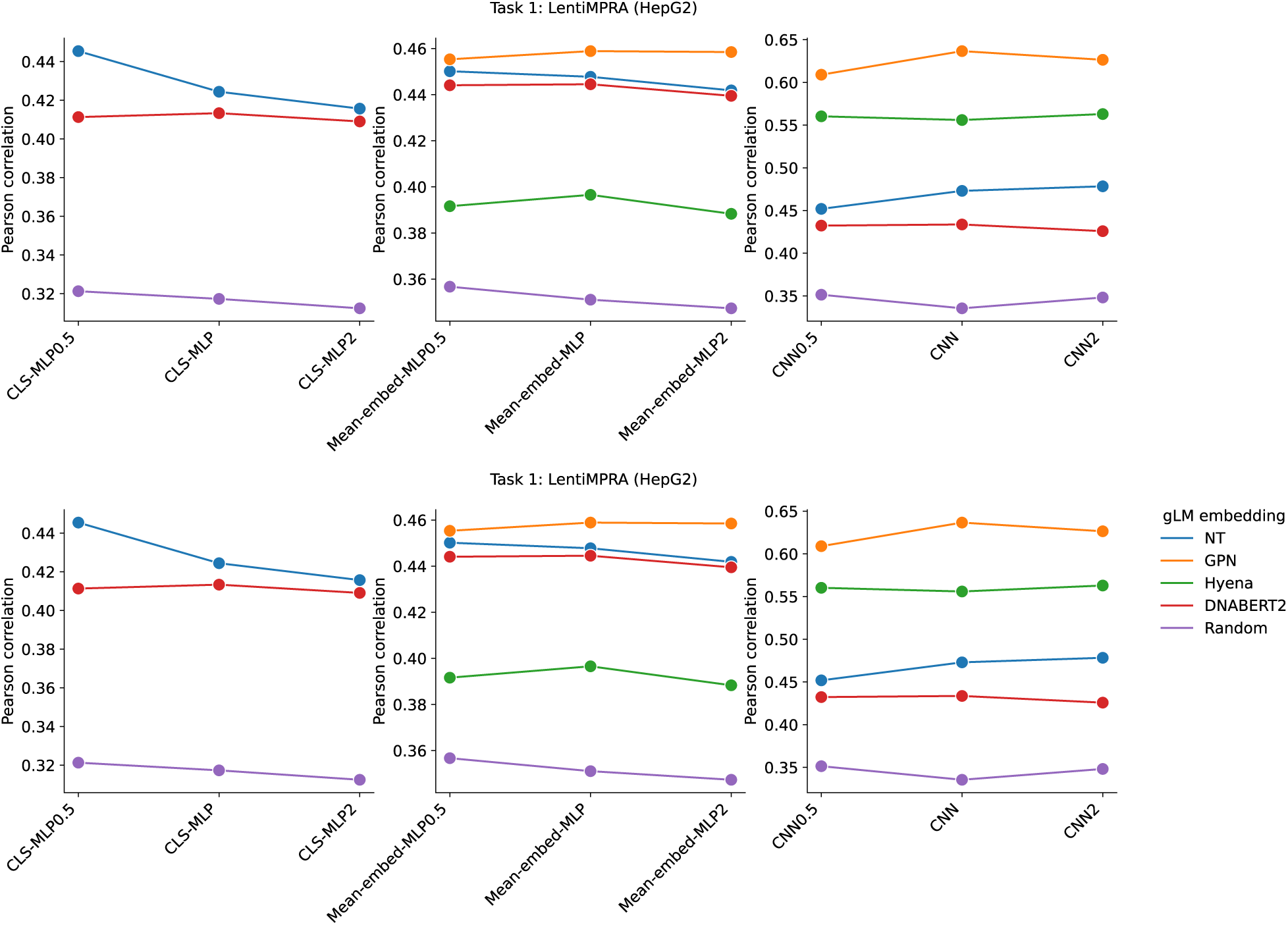
Comparison of performance for various hyperparameter choices for downstream models on lentiMPRA data. Four types of downstream models, including ridge regression (Ridge), multi-layer perceptron (MLP), and convolutional neural network (CNN), were trained for each gLM. The numbers after the model names (i.e., 0.5 or 2) represent the scaling factor for the number of parameters in each hidden layer relative relative to the original model structure. These results show that, up to a factor of 2, the performance stays roughly similar.

**Supplementary Figure 2.**
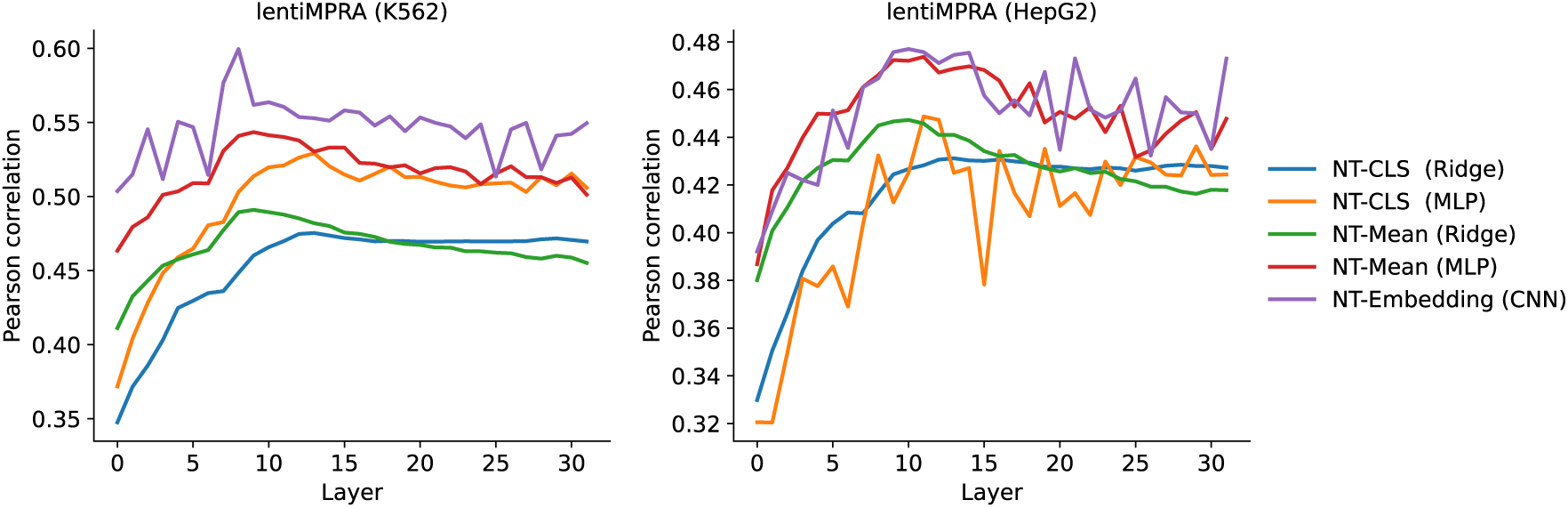
Layer-wise performance of Nucleotide-Transformer on the lentiMPRA dataset. Test performance of various machine learning models trained using embeddings from different layers of Nucleotide-Transformer. Embeddings include the CLS token, mean embedding (Mean), and the full embedding (Embedding). Machine learning models include ridge regression (ridge), multi-layer perceptron (MLP) and a convolutional neural network (CNN).

**Supplementary Figure 3.**
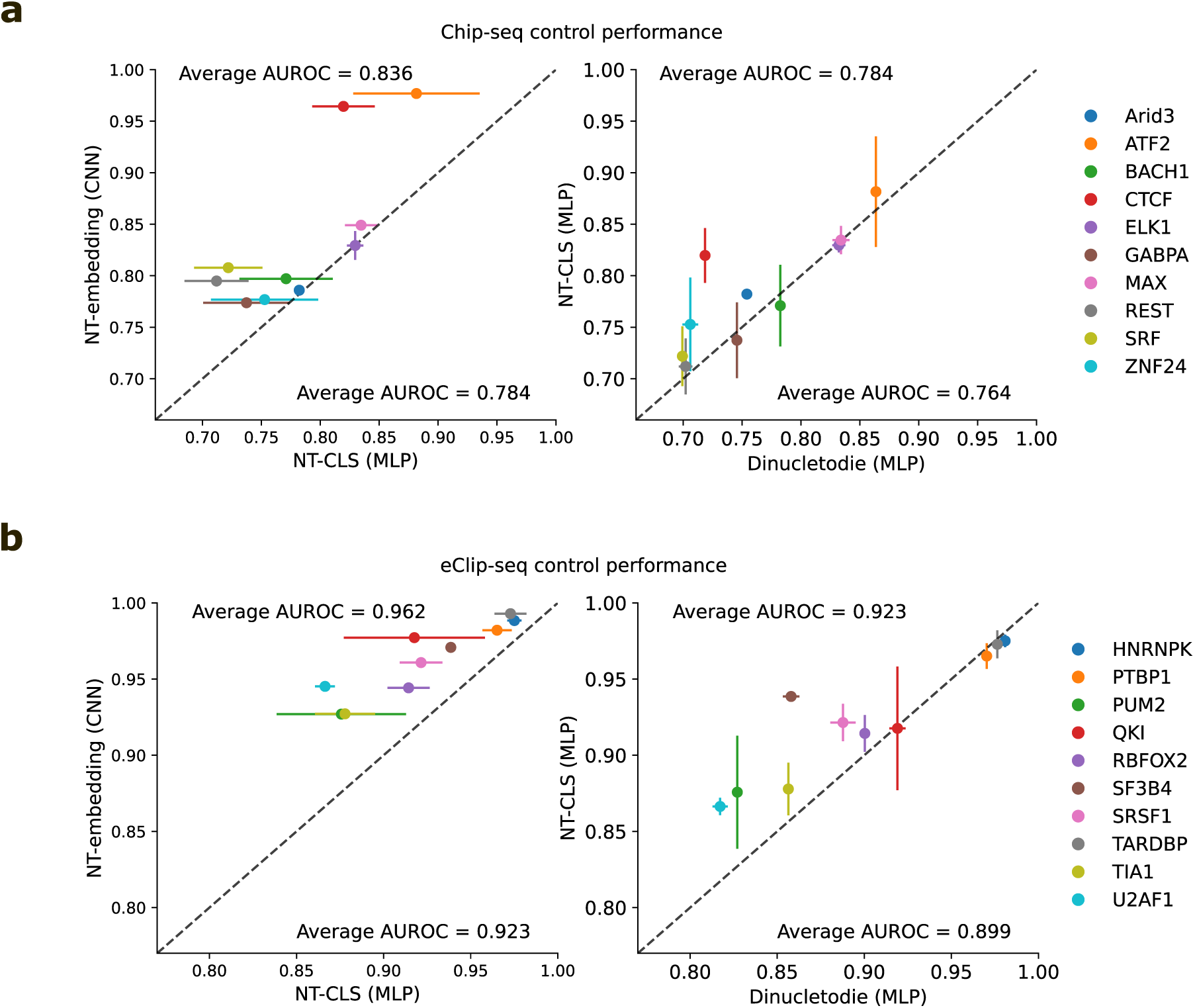
Control experiments with different embeddings. Performance comparison between a CNN trained using full embeddings of the penultimate layer from Nucleotide-Transformer, an MLP trained using Nucleotide-Transformer’s CLS token, and an MLP trained using dinucleotide frequencies of the sequence on (**a**) ChIP-seq data and (**b**) eCLIP-seq data. Performance is measured by the average area-under the receiver-operating characteristic curve (AUROC) and error bars represent the standard deviation of the mean across 5 different random initializations. Text valeus represent the average AUROC across all ChIP-seq or CLIP-seq datasets.

**Supplementary Figure 4.**
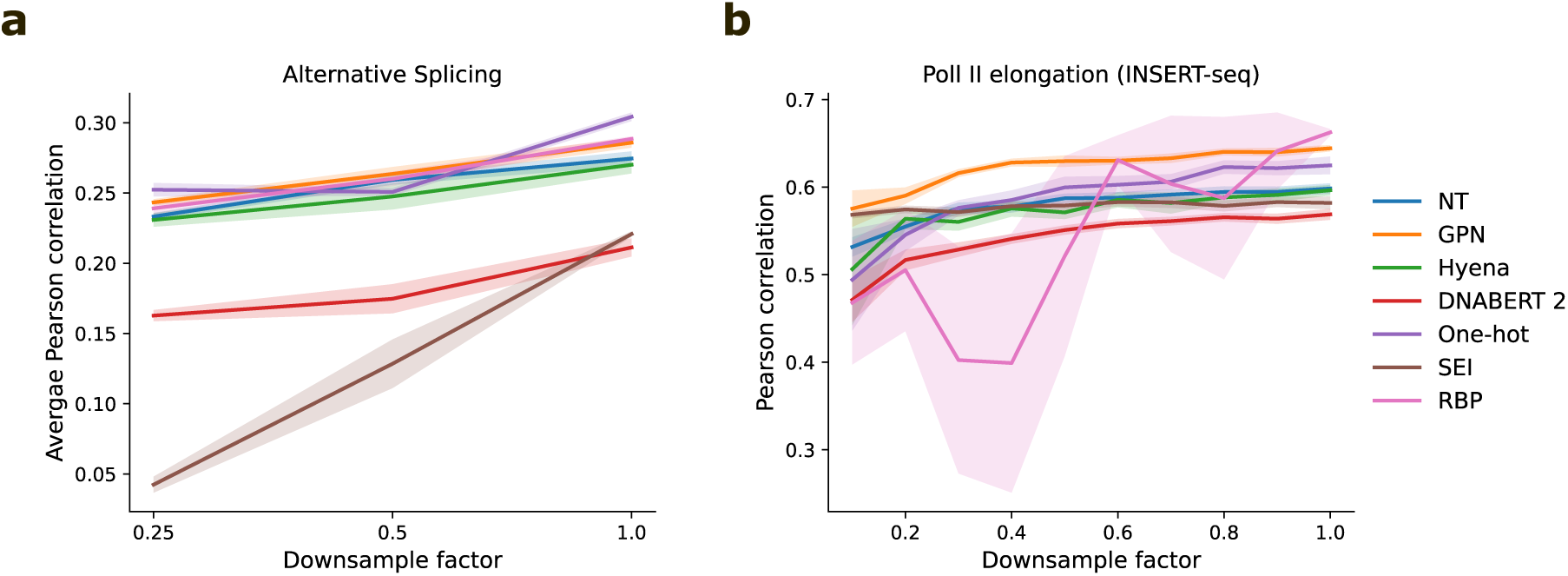
Down-sampling performance on RNA regulation tasks. Average performance of machine learning models on (**a**) alternative splicing, task 4, and (**b**) RNA Pol II elongation potential, task 5, down-sampled by various factors. Shaded region represents standard deviation of the mean across 5 different random initializations.

**Supplementary Figure 5.**
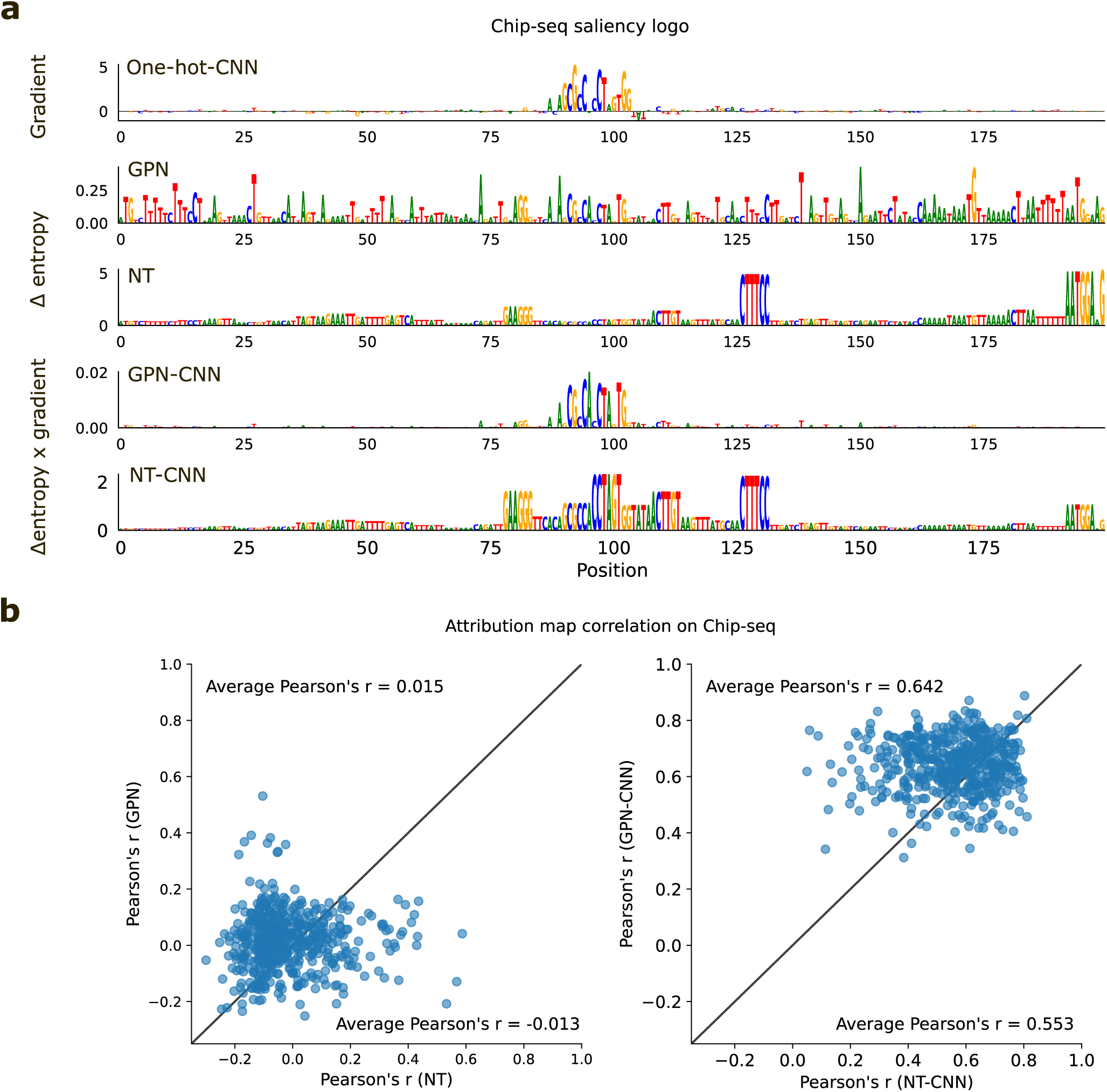
Attribution analysis comparison for sequences from CTCF ChIP-seq data. **a**, Representative example of attribution maps for a CTCF binding sequence. Attribution maps include (top to bottom): the gradient-times-input of a one-hot-trained CNN; the delta entropy of predicted nucleotides via single-nucleotide masking from a pre-trained GPN; the delta entropy of predicted nucleotides via single-nucleotide masking from a pre-trained Nucleotide-Transformer; the gradient of a CNN-trained using GPN embeddings multiplied by the delta entropy of predicted nucleotides via single-nucleotide masking from a pre-trained GPN; and the gradient of a CNN-trained using Nucleotide-Transformer embeddings multiplied by the delta entropy of predicted nucleotides via single-nucleotide masking from a pre-trained Nucleotide-Transformer. **b**, Scatter plot comparison of the attribution map correlations for different pre-trained gLMs (left) and CNNs trained using gLM embeddings (right). Attribution map correlations reflect the Pearson correlation coefficient between the attribution map generated by the gLM-based attribution method with the Saliency Map generated by a one-hot-trained CNN. Each dot represents a different sequence in the CTCF ChIP-seq dataset (N=500).

**Supplementary Figure 6.**
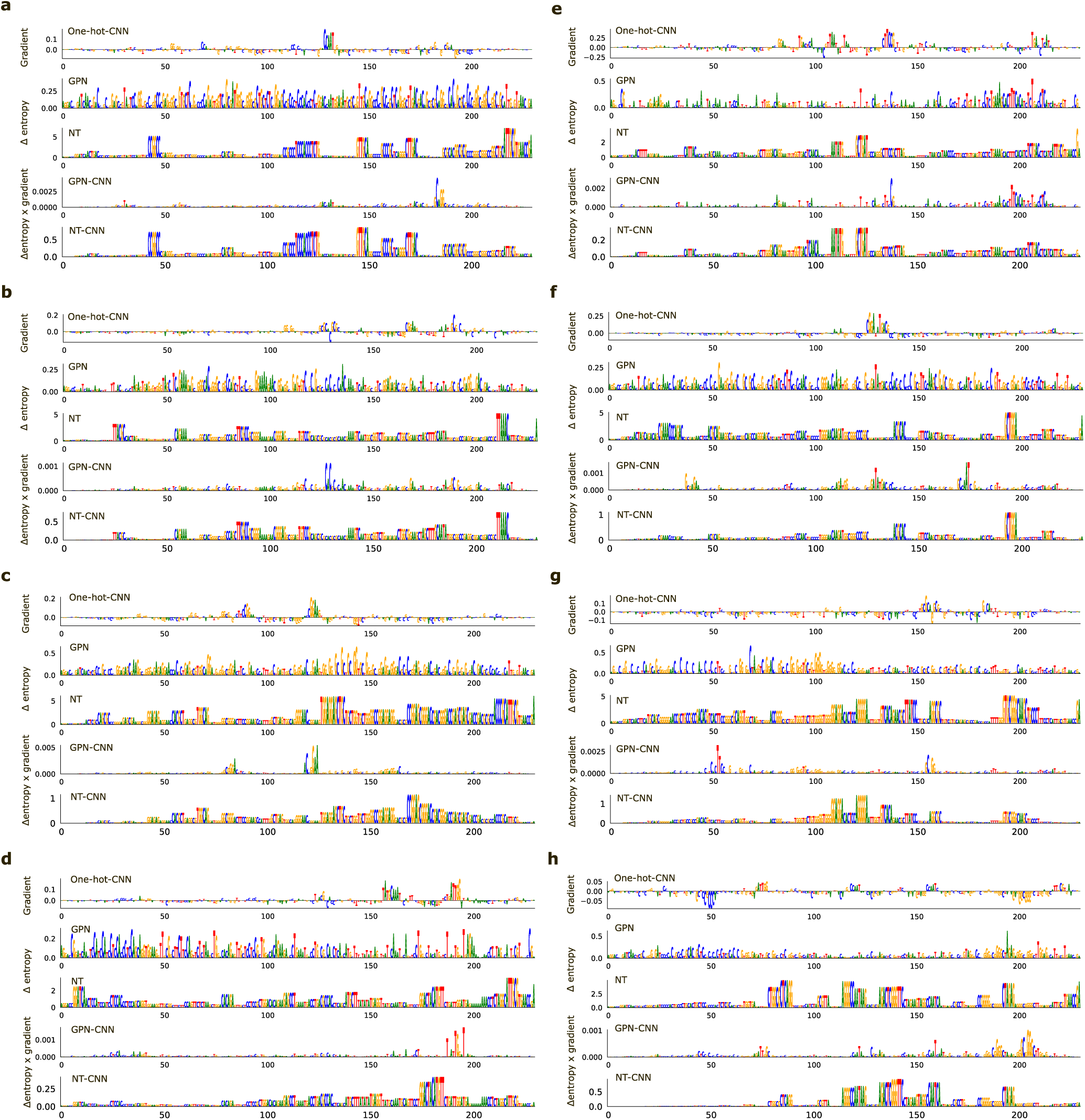
Representative examples of attribution maps for sequences from the lentiMPRA dataset. In each panel, attribution maps are shown for different sequences in order of (top to bottom): the gradient-times-input of a one-hot-trained CNN; the delta entropy of predicted nucleotides via single-nucleotide masking from a pre-trained GPN; the delta entropy of predicted nucleotides via single-nucleotide masking from a pre-trained Nucleotide-Transformer; the gradient of a CNN-trained using GPN embeddings multiplied by the delta entropy of predicted nucleotides via single-nucleotide masking from a pre-trained GPN; and the gradient of a CNN-trained using Nucleotide-Transformer embeddings multiplied by the delta entropy of predicted nucleotides via single-nucleotide masking from a pre-trained Nucleotide-Transformer.

## Notes

### Competing Interest Statement

The authors have declared no competing interest.

### Summary of Updates

Updated analyses (included in supplementary Tables 2 and 3) and minor updates to the main text.

